# The role of uncertainty and negative feedback loops in the evolution of induced immune defenses

**DOI:** 10.1101/2022.10.11.511650

**Authors:** Danial Asgari, Alexander J. Stewart, Richard P. Meisel

## Abstract

Organisms use constitutive or induced defenses against pathogens, parasites, and herbivores. Constitutive defenses are constantly on, whereas induced defenses are activated upon exposure to an enemy. Constitutive and induced defenses each have costs and benefits, which can affect the type of defense that evolves in response to pathogens. Previous models that compared the two lacked mechanistic details about host defense, did not consider pathogen proliferation rates, or lacked both features. To address this gap, we developed a detailed mechanistic model of the well-characterized *Drosophila melanogaster* immune signaling network. We evaluated the factors favoring the evolution of constitutive and induced defenses by comparing the fitness of each strategy under stochastic fly-bacteria interactions. We show that an induced defense is favored when bacteria are at low density, heterogeneously distributed, or have fluctuating distributions in ways that depend on the bacterial proliferation rate. Our model also predicts that the specific negative regulators that optimize the induced response depend on the bacterial proliferation rate. We therefore conclude that an induced immune defense is favored in environments where bacterial encounters are uncertain—because of heterogeneity in spatial or temporal distributions—but that benefit depends on the mechanism of induction and pathogen properties.

## Introduction

Organisms defend themselves against natural enemies—including pathogens, parasites, and herbivores—using induced and constitutive defenses. Induced responses are produced when an enemy is present or an organism is attacked, whereas constitutive defenses are always on. Induced and constitutive defenses each offer distinct advantages, yet also have limitations. For example, constitutive strategies may be costly because of the resources required to produce the defenses or direct deleterious (immunopathological) effects of defenses on host tissues (Aggarwal & Silverman, 2008; Li et al., 2020). It may be more cost effective for an organism to deploy an induced strategy in which defensive molecules are only produced when needed, such as when the host is attacked by an enemy (Karban, 2020). However, this induced strategy can create a costly delay in response (Karban & Myers, 1989), although priming might reduce this delay (Pham et al., 2007). If the cost of defense is low or delay in response is too costly, a constitutive strategy might therefore be favored (Bixenmann et al., 2016).

Mathematical models have revealed insights into the evolution of induced and constitutive defenses. For example, Clark and Harvell (1992) used a minimal model to study resource allocation to growth, reproduction, and defense. They showed that an induced defense is favored when attacks are unpredictable. Later models that solely focused on the cost of defense found that increasing the probability of attacks favors a constitutive defense (Kamiya et al., 2016; Shudo & Iwasa, 2001). Other theoretical models have explored how different degrees of uncertainty shape the evolution of constitutive and induced defenses. For example, Adler and Korban (1994) showed that, in the absence of environmental variation (i.e., there is certainty of an attack), a constitutive defense is favored. On the other hand, Hamilton et al. (2008) found that uncertainty about the parasite proliferation rate favors induction. However, none of this work considered the actual biological mechanisms underlying induced and constitutive defenses, including the signaling pathways responsible for induction.

The well-characterized innate immune response of *Drosophila melanogaster* is a good model system for incorporating biological mechanisms into our understanding of the evolution of induced and constitutive defensive strategies. *D. melanogaster* uses a combination of constitutive and induced expression of antimicrobial peptides (AMPs), along with other effectors, to fight off pathogens (Lemaitre & Hoffmann, 2007). Induced and constitutive expression of AMPs is under tissue-specific control in *D. melanogaster.* For example, the constitutive expression of AMPs in the *D. melanogaster* salivary gland and ejaculatory duct is under the control of a single transcription factor, Caudal (Ryu et al., 2004).

The *Drosophila* induced immune response depends on the microbial pathogen, as well as which tissue is infected. The immune deficiency (Imd) and Toll pathways are the two major signaling cascades involved in the *Drosophila* induced immune response to bacterial and fungal infection (De Gregorio et al., 2002). The Toll pathway regulates the expression of AMPs upon exposure to glucans found in fungal cell walls or lysine-type peptidoglycans (PG) present in the cell wall of Gram-positive bacteria (El Chamy et al., 2008). The Imd pathway is activated by diaminopimelic acid (DAP)-type PG in the cell walls of Gram-negative bacteria (Kaneko et al., 2005). The Imd pathway is active in the fat body and gut, which are the primary immune organs of flies (Myllymäki et al., 2014). Following ingestion, bacteria proliferate inside the gut and release PG, which activates the Imd pathway (Neyen et al., 2012). The Toll pathway, in contrast, is not active in the gut because it relies on cytokines for activation, which are not stable in the low pH environment of the digestive tract (Buchon et al., 2009; Ryu et al., 2006). Because flies in nature are infected by bacterial pathogens mainly by ingestion during feeding (Siva-Jothy et al., 2018), modeling the Imd pathway in response to bacteria is more ecologically relevant.

We developed a mathematical model of the well-characterized *D. melanogaster* Imd pathway (Figure 1). The first step of Imd pathway activation entails binding of bacterial PG to receptor proteins (PGRP-LC and -LE) on the surface of *Drosophila* enterocytes (Kaneko et al., 2006; Schmidt et al., 2008). Next, a signaling complex—consisting of the Imd protein, the adaptor protein dFADD, and the caspase protein DREDD—is recruited to the intracellular domain of the receptor (Georgel et al., 2001; Leulier et al., 2002). DREDD cleaves Relish, which is an NF-κB transcription factor (Stoven et al., 2003), and the N-terminal domain of Relish is translocated into the nucleus where it facilitates the transcription of AMP genes (Stöven et al., 2000). Imd signaling concurrently activates negative feedback loops, which protect the host from harmful effects of excessive immune response (Badinloo et al., 2018). For example, scavengers of PG (PGRP-LB) are expressed in a Relish-dependent manner, which reduce Imd signaling by removing PGs from the gut (De Gregorio et al., 2002). Relish also upregulates *pirk*, which encodes a negative regulator of the Imd pathway. Pirk reduces the number of available PGRP-LC receptors on the cell surface (Kleino et al., 2008; Lhocine et al., 2008). Finally, following activation of the Imd pathway, two transcription factors, Jra and Stat92Ea, form a repressosome complex with a high mobility group protein DSP1. The repressosome competes with Relish over binding to promoters of AMP genes, thereby reducing the AMP expression (Kim et al., 2007). The production of the transcription factor Jra has been shown to be regulated by Relish, thus acting as a negative feedback loop to attenuate Imd signaling (De Gregorio et al., 2002).

**Figure 1.**
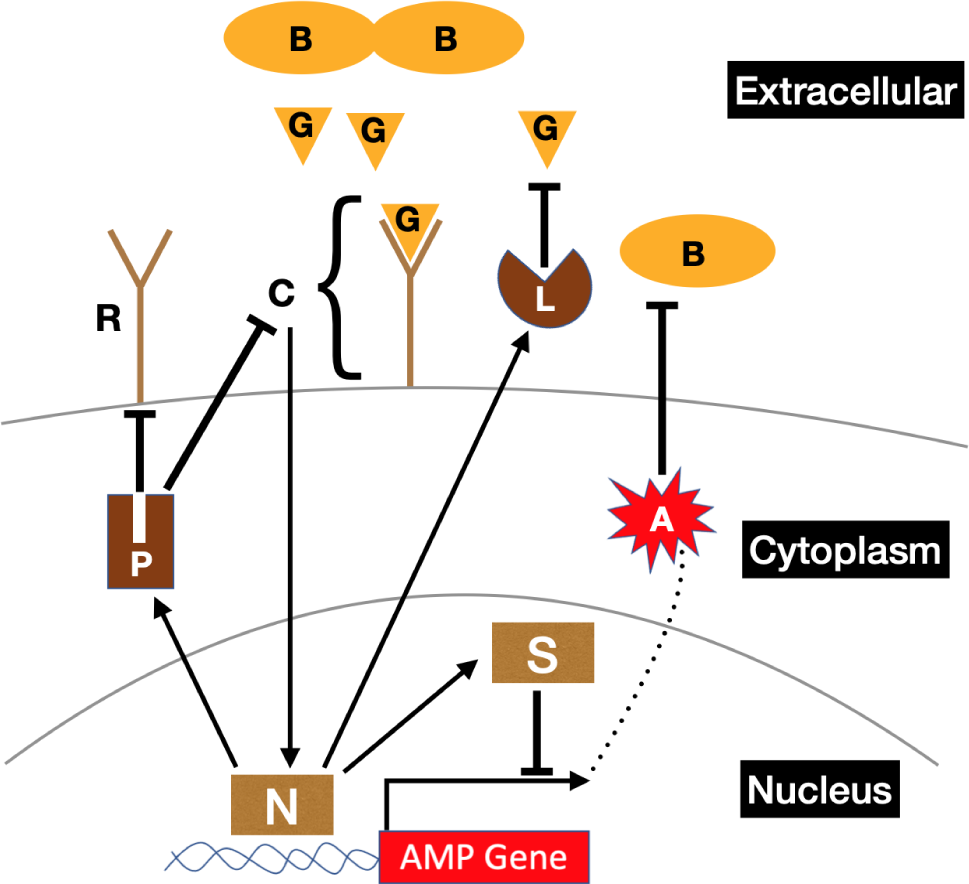
Diagram showing the Imd signaling pathway for induction of antimicrobial peptides (A). The letters in the diagram correspond to the variables in the system of ordinary differential equations of the induced model (Eqs. 1-9). Peptidoglycans (G) produced by bacteria (B) bind to cell surface receptors (R), forming a receptor complex (C) that initiates an NF-κB signaling pathway to activate the transcription factor Relish (N). Relish promotes the transcription of AMP genes, which leads to the production of AMPs (A). Relish also induces the production of Pirk (P), scavengers of peptidoglycans (L), and at least one gene encoding a component of the repressosome complex (S). AMPs destroy bacteria, and Pirk reduces available cell surface receptors. Repressesome (S) competes with Relish (N) for binding to the promoter of AMP genes.

To investigate the conditions that favor induced versus constitutive defenses, we used a system of ordinary differential equations (ODEs) to model the Imd response to bacteria in *D. melanogaster*. Our model of the Imd signaling pathway goes beyond one previously developed by Ellner et al. (2021) because ours includes mechanistic details of negative feedback loops involved in the pathway. We used this detailed model to explore how negative regulators modulating the immune response at different steps in the signaling pathways that control induction shape the fitness of an induced defense when the host encounters different bacterial populations. We compared the fitness of this model to a simple model of constitutive defense. We did not consider mixed strategies as it has been shown that immune genes can adopt a purely constitutive or a purely induced strategy (Asgari et al., 2022). Both our induced and constitutive models include features that capture the bacterial proliferation rate and survival in the gut, which has been shown experimentally to vary across bacteria species (Duneau et al., 2017; Pais et al., 2018). We considered stochastic encounters of flies with bacteria in a variety of environments, with different bacterial density and patchiness, to determine the conditions in which induced or constitutive defenses are favored.

## Materials and Methods

### The model

We modeled the Imd pathway using a system of ODEs which capture the dynamics of immune response to bacterial infection (Figure 1). Equation (1) describes the change in bacterial density (*B*) in the gut of an individual fly.

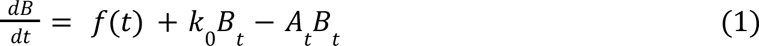

Under our model, the density of bacteria entering the gut at time *t* is described by the function *f*(*t*). In general, *f*(*t*) depends on the local concentration of bacteria encountered by the fly as it moves around its environment. For example, we may consider scenarios in which *f*(*t*) is a deterministic, oscillating (sinusoidal) function, reflecting an environment in which bacteria occur in regular “patches”. We also consider scenarios in which *f*(*t*) describes the probability of entering or leaving a patch in an environment with randomly distributed colonies of bacteria. After ingestion, bacteria proliferate inside the gut at rate *k*_0_, and are killed by AMPs at a rate *A_t_ B_t_*, where *A_t_* is the concentration of AMP at time *t*. In order to contrast the Imd pathway with a constitutive immune response, we modeled constitutive expression by simply assuming that *A_t_* is a constant (*A_t_* = *A*) in Equation (1). The model for the constitutive defense only contains Equation 1 because signaling proteins are not involved in constitutive defenses.

The concentration of free bacterial PG at time *t,* (*G_t_*), is described by Equation (2).

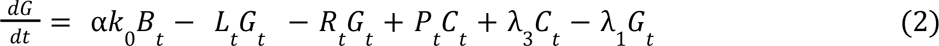

The production of free PG is stimulated by the proliferation of bacteria (*k*_0_ *B_t_*) at rate α. We assume that free PG is scavenged (i.e., removed from the system) by PGRP-LB (*L_t_*) interacting with *G_t_* (Costechareyre et al., 2016; Zaidman-Rémy et al., 2006). Free PG is also lost when it binds to receptors on the cell surface (*R_t_*). We also assume that PG is released when the PG-receptor complex (produced at a rate *C_t_*) is pulled down by Pirk (produced at a rate *P_t_*) (Kleino et al., 2008). The receptor-PG complex is dissociated at rate λ_3_, which produces free PG and free receptors. Finally, PG is degraded at rate λ_1_.

Next, we describe the change in concentration of free receptor (*R*) over time via Equation (3).

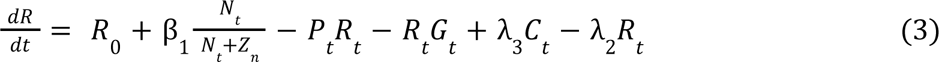

Here, receptors are produced at a base rate *R*_0_. Receptor expression is positively regulated by the NF-κB transcription factor Relish (*N*_*t*_) at rate β_1_, which is modeled as a Hill function (Liu et al., 2020). *Z*_*n*_ is the binding energy of Relish to the promoter, which is inversely proportional to the probability of binding. Interactions with Pirk and PG reduce the concentration of free receptors. We considered both separate and identical degradation rates for *R*, *N*, *L*, *P*, *S*, and *A* (described below). Equations in the main text have equal degradation rates for *R*, *N*, *L*, *P*, *S*, and *A* (λ_2_).

Following from Equation (3), change in the concentration of receptor-PG complex (*C*) is described by Equation (4).

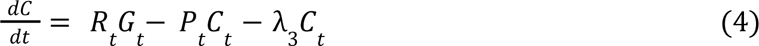

Equation (5) describes the change in the NF-κB transcription factor Relish, which is activated at rate β_2_ following formation of the PG-receptor complex (Stoven et al., 2003).

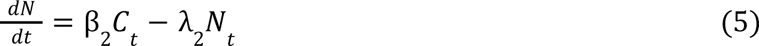

Equations (6), (7), and (8) describe negative feedback loops mediated by Relish.

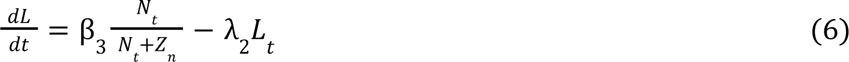

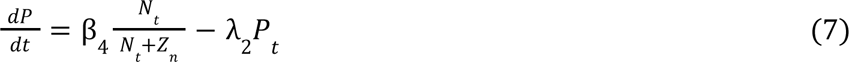

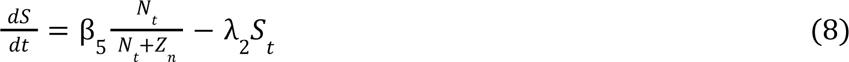

PGRP-LB, Pirk, and the repressosome (AP-1 and STAT complex, *S_t_*) are activated by Relish at rates β_3_, β_4_, and β_5_, respectively (Kim et al., 2007; Kleino et al., 2008; Lhocine et al., 2008; Zaidman-Rémy et al., 2006). Equation (9) describes AMP expression, which is upregulated by Relish and repressed by the repressosome with a single rate of β_6_. The repressosome competes with Relish for binding to promoters of AMPs (Kim et al., 2007). The binding energy for the repressosome is *Z_s_*.

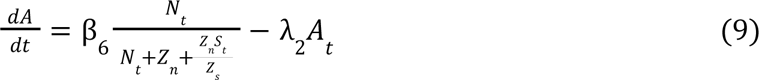

### Simulation of the bacterial environment

In order to capture the effect of stochastic interactions with bacterial colonies on immune response, we simulated bacterial populations on a 2-dimensional lattice (100 × 100) using a simple random walk algorithm with periodic boundary conditions (Figure 2). We placed bacterial colonies at positions on the lattice which were determined according to a random walk. We initialized the walk at a random starting coordinate [*i*_0_, *j*_0_], and the subsequent direction of movement was specified to be left [*i*_0_ − *p*, *j*_0_], right [*i*_0_+ *p*, *j*_0_], up [*i*_0_, *j*_0_ + *p*], or down [*i*_0_, *j*_0_− *p*], with equal probabilities 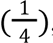, where *p* describes the (constant) step size. A bacterial colony was then placed on the lattice at the updated coordinate [*i*, *j*]. If a lattice point was already occupied by a bacterial colony, the lattice was not updated, the colony count remained unchanged, and another step was taken. The process was repeated until a predetermined number of colonies had been added to the lattice. The number of colonies added to the lattice determined the density of bacteria 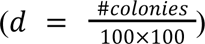 in the environment. The value of the random walk step size, *p*, determines the patchiness of the bacterial distribution; low values of *p* create a more heterogeneous distribution of bacteria, while high values create uniform distributions (Figure 2).

**Figure 2.**
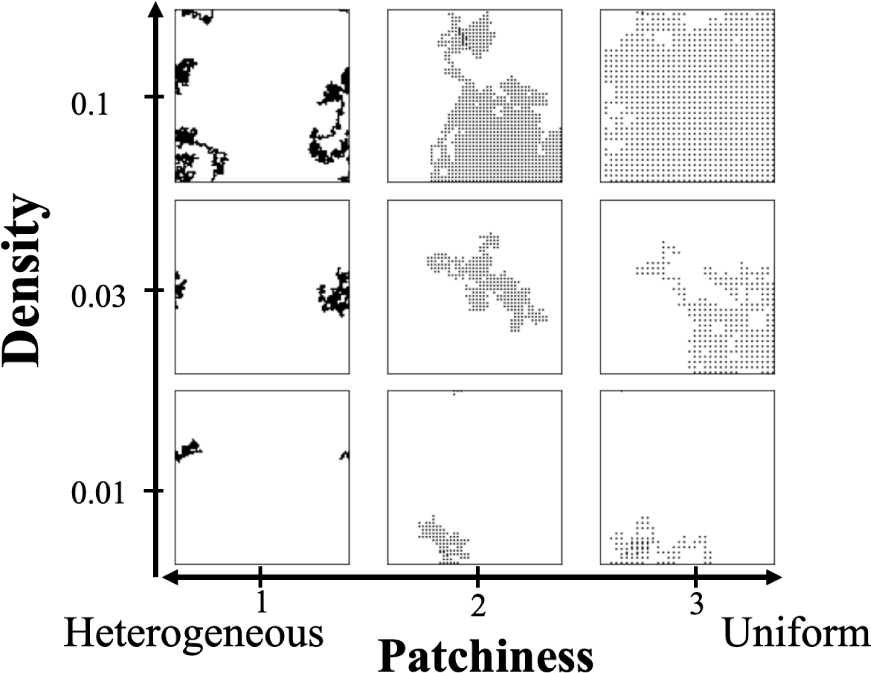
Examples of simulated environments with different numbers (Density) and patchiness of bacterial colonies. Each dot represents a bacterial colony on a 100 × 100 lattice.

### Movement of the fly through the environment

We used both a deterministic and stochastic approach to model fly movement through bacterial environments. In the deterministic approach, we considered an oscillating model in which *f*(*t*) = ω *sin*(*t* Φ)^2^, such that the fly experiences a deterministic oscillating bacterial environment. The frequency of encounter is captured by Φ, and the amount of bacteria is represented by ω. For the deterministic oscillating model, we solved the system of ODEs using the scipy python package.

In the stochastic method, we assessed the efficacy of constitutive defense and induced immune response by comparing their efficacy over the course of a large number of sojourns through simulated bacterial environments (Figure 2). To those ends, we simulated the behavior of a fly within a given bacterial environment as a random walk across a lattice seeded with a bacterial distribution as described above. The random walks were performed as described above for seeding the lattice, but all fly walks used step size *p* =1. When a fly landed at a given coordinate [i,j], the immune response was determined by the input function *f*(*t*) (Eq. 1), such that, if a bacterial colony was present *f*(*t*)=1, and *f*(*t*)=0 otherwise. Thus as the fly moves through the environment *f*(*t*) takes values 1 or 0 depending on the presence or absence of bacteria at the current lattice position. This process simulates stochastic fly-bacteria interactions. We solved the system of ODEs using the Runge-Kutta Method (RK4) algorithm for the random walk model.

Two parameters in our induced model define the interaction between the bacteria that flies encounter and their immune system during their deterministic oscillations and random walks: the rate at which PG is released upon bacterial proliferation (α) and the degradation rate of PG (λ_1_). We considered four values for α (0.2, 1, 2, 4) and two values for λ_1_ (0.01 and 0.05). However, our results focus on λ_1_ = 0.01 for reasons described in the Results. In both the induced and constitutive models, the efficacy of the immune response also depends on the bacterial proliferation rate (*k*_0_). We tested both constitutive and induced defenses against three different bacterial proliferation rates (*k*_0_= 0.1, *k*_0_= 0.2, and *k*_0_= 0.5).

### Calculating fly fitness

After obtaining a vector containing values at every time point for each variable in the model (*B_t_*, *G_t_*, *R_t_*, *C_t_*, *N_t_*, *L_t_*, *P_t_*, *S_t_*, *A_t_*), we calculated fitness as an exponentially decreasing function of the summed time averaged values of each of the model variables, as shown in Equation (10) for an induced response and in Equation (11) for constitutive defenses.

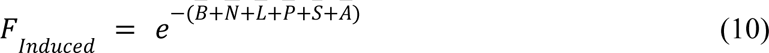

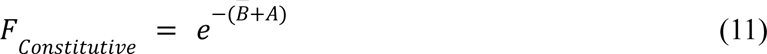

In these calculations (Eqs. 10–11), 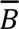 is the arithmetic time averaged bacterial concentration in the gut, 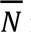 is the time averaged production of Relish, and so on. This method assumes that fitness declines exponentially with the average bacterial load 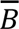 and with the strength of the immune response due to the cost of production. We explored the effect of the fitness function on our conclusions, as described in the Results.

For each environment with a particular bacterial density and patchiness, we simulated 1,000 fly random walks each consisting of 100,000 steps, and we calculated the arithmetic mean fitness for the constitutive and induced defenses across simulations. The fitness difference (Δ*F*) between the induced and constitutive immune responses was calculated by subtracting the average fitness of the induced response 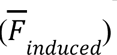 from the average fitness for the constitutive defense 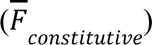 using Equation (12).

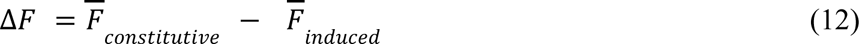

### Parameter optimization

The model for induced response has 11 parameters that describe the attributes of the Imd pathway (λ_2_, λ_3_, *R*_0_, β_1_, β_2_, β_3_, β_4_, β_5_, β_6_, *Z_n_*, *Z_s_*). To find the parameters that maximize the fitness of the induced response, we simulated random walks within the 11-dimensional landscape consisting of the 11 parameters, which we refer to as “optimization walks”. We used the Muller (1959) method to generate random points in an 11-dimensional space, where step size = 0.01. First, we found the position vector of a point (*u*) in an 11-dimensional space by generating 11 independent normal deviates (Eq. 13).

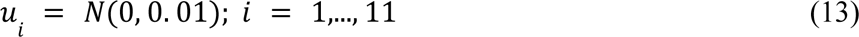

The length of the vector is calculated using Equation (14).

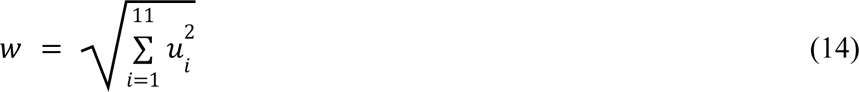

The updated value for each variable (*X*’) is found by summing the current value (*X*) with the direction cosines 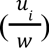 multiplied by the step size (0.01), using Equation (15).

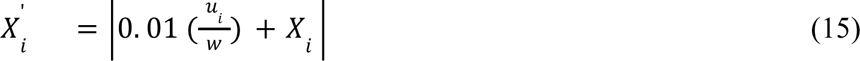

We used Equation (10) to calculate the fitness of induced defense (*F_induced_*) using the 11 parameters as input. At each step in the optimization walk, the 11 parameters are simultaneously updated. If 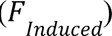, the new set of parameter values is accepted; otherwise, the set is rejected. We used 10,000 steps for the lower bacterial proliferation rates (*k*_0_ = 0.1 and *k*_0_ = 0.2), and 20,000 steps for the higher proliferation rate (*k*_0_ = 0.5) because it took longer for fitness to plateau for the higher proliferation rate.

Random walks within an 11-dimensional landscape are computationally intensive if *f*(*t*) is stochastic. To perform a more efficient search, we calculated fitness at each step of our random walk using a deterministic oscillating input with different frequencies to the system of ODEs instead of a stochastic one(*f*(*t*) = *sin*(*t* Φ)^2^).. We optimized the induced model using different values of Φ, i.e., the frequency of the sinusoidal input. Higher Φ describes a higher density of bacteria. Using a sinusoidal input is more efficient than stochastic simulations of bacterial distributions at each step of the walk through parameter space, and it also dispenses with the need for simulating random walks of flies among bacteria at each step. Different local optima might be reached when starting from different points in the landscape. Therefore, we considered a large and a small starting value for each parameter and ran a random walk for every combination of large and small parameter values. This makes a total of 2048 walks for the purpose of optimizing the 11-parameter model (2^11^ = 2048).

To test for the robustness of the assumption of identical degradation rates (λ_2_) in the 11-parameter model, we also considered a model with different degradation rate parameters for *R*, *N*, *L*, *P*, *S*, and *A*. This created a 16-parameter model. Optimizing this model requires a total of 65536 (2^16^) random walks, which is 5 orders of magnitude more computationally intensive than the 11-parameter model. To overcome this problem, we optimized the 16-parameter model using two approaches. In the first approach, we considered a large and a small starting value for the six degradation rates and started the other 10 parameters at their optimum values in the 11-parameter model. In the second approach, we also considered a large and a small starting value for the six degradation rates but fixed the other 10 parameters at their optimum values in the 11-parameter model. Both approaches entail a total of 64 (2^6^) walks for the purpose of optimizing the 16-parameter model, following optimization of the 11-parameter model.

To optimize the constitutive defense, the amount of constitutively expressed AMP (*A*) was changed over a range of 0.01 to 2 (step size = 0.01), and the value of *A* that confers the highest fitness is chosen. The Euler method was used (h = 0.01) to solve Equation (1), where *A_t_* is a constant (*A _t_*= *A*). Due to the simplicity of the model for constitutive defense, the model was optimized in stochastic environments. The optimum fitness for the constitutive defense was calculated in each combination of bacterial density, patchiness, and proliferation rate and then compared to the fitness of different optimizations of the induced defense (i.e., induced defenses optimized with different values of Φ). The induced and constitutive defenses that are tested against each other are optimized using the same bacterial proliferation rates.

### Simulating fluctuation across multiple environments

We simulated fluctuation in density and/or patchiness of bacterial populations. To those ends, we randomly selected *j* environments with different densities and/or patchiness of bacteria, and we assumed that flies encounter those environments with equal probabilities. Sampling was done without replacement in order to maximize fluctuations in the density and/or patchiness of the bacterial population. To estimate the fitness of flies encountering *j* bacterial environments with equal probabilities, we calculated the arithmetic mean of the fitness values across *j* environments (Eq. 16).

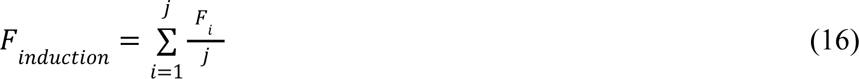

In each of the *j* environments, the fly stochastically encounters bacteria. Simulations were performed for induced defenses that were optimized with different values of Φ.

To find the optimum fitness for the constitutive defense when flies inhabit *j* environments with equal probabilities (*j*^−1^), we calculated the fitness in each environment for constitutive defenses ranging from 0.01 to 2 units of AMP production (i.e., 0.01 ≤ *A* ≤ 2). Next, we calculated arithmetic means of fitness values across *j* environments (the dashed lines in supplemental figure 1) and chose the maximum value (the arrows in supplemental figure 1). We measured the proportion of times induction outperforms constitutive defense by repeating the process of random selection of environments and calculation of the relative fitness (Δ*F*) 10,000 times.

We also calculated the relative fitness of the constitutive and induced defenses when the fly inhabits 2 environments with probabilities *q* and 1 − *q*. This differs from the previous analysis, where *q* = *j*^−1^. The fitness of the induced response is the weighted average of optimum fitness values across the two environments (Eq. 17).

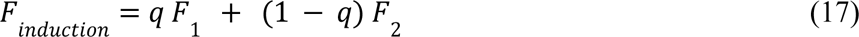

We used the following approach to find the optimum value of AMP production (*A*) for the constitutive defense when flies inhabit two environments with probabilities *q* and 1 − *q*. First fitness was calculated in each of the two environments for constitutive defenses ranging from 0.01 to 2 units of AMP production (0.01 ≤ *A* ≤ 2). The optimum fitness of constitutive defense is the maximum value (arrows in supplemental figure 2) of the weighted averages of fitness values (dashed lines in supplemental figure 2). The proportion of times induction outperformed constitutive defense for 10,000 combinations of environments was calculated for each value of *q*, as explained above.

## Results

### Induction is favored when bacteria are at low/intermediate density and have heterogeneous distributions

In order to compare the fitness of induced and constitutive defenses, first we found parameter values in our model that maximize fitness for each strategy in a variety of environments. Constitutive defense was optimized by finding the value of constitutively expressed AMP (*A*) that produces the highest fitness within each environment, where the environment consists of a specified density (*d*) and patchiness (*p*) of bacteria. To optimize the induced response, we simulated random walks in an 11-dimensional landscape corresponding to the 11 parameters in our model, and we identified the combination of 11 parameter values that maximize fitness in a specified environment. We optimized the induced response across a variety of environments with different frequencies of bacterial encounters using a sinusoidal function, where Φ corresponds to the frequency of bacterial exposure (optimized parameter values are reported in supplemental table 1). Induced defenses are plastic defenses that evolve when environmental fluctuations are predictable (Scheiner, 1993). Therefore, optimizing the induced defense by a sinusoidal function, which simulates predictable fluctuations, is a valid approach for producing effective induced immune defenses. Higher values of Φ correspond to environments with a higher density and more uniform distribution of bacteria, where fly-bacteria interactions are more likely. Lower values of Φ, in comparison, correspond to environments with low density or more heterogeneous bacterial distributions, where bacterial exposure is lower.

We evaluated how the rate of bacterial PG production affects the performance of an induced immune response across different frequencies of bacterial exposure. We optimized induced defenses for four rates of bacterial PG production (α = 0.2, 1, 2, and 4; supplemental figure 3A). Our results were qualitatively similar across different α and Φ values (supplemental figure 3B). Induction performs best when bacterial density is intermediate and the distribution is heterogeneous, regardless of the α and Φ values. For the remainder of the manuscript, we only focus on results for α = 2 because it maximizes the performance of induced defenses when flies encounter two different bacterial distributions (supplemental figure 3C).

We chose a small value for the natural degradation rate of PG (λ_1_= 0.01) because the natural degradation rate is much slower compared to degradation by PGRPs (Filipe et al., 2005). We found that, when λ_1_= 0.01, the parameter controlling production of PGRP-LB (β_3_) is larger than λ_1_ (supplemental figure 4A), consistent with more degradation by PGRP-LB than the natural degradation rate. However, when λ_1_ = 0.05, β_3_ is smaller (supplemental figure 4B). Therefore, setting λ_1_to 0.01 is biologically realistic because it captures the important role that PGRP-LB plays in degradation of PG. By setting α = 2 and λ_1_= 0.01, we found that increasing the frequency of fly-bacteria encounters (Φ) reduced the optimal fitness of the induced response, regardless of the bacterial proliferation rate (*k*_0_) inside the fly (supplemental figure 5). Our results therefore suggest that induced responses are less fit in environments with a higher frequency of fly-bacteria interactions.

We compared the fitness of the induced and constitutive defenses under a sinusoidal mode of encounter with bacteria (*f*(*t*) = ω *sin*(*t* Φ)^2^).. For most optimizations of the induced defense, an induced response is favored for at least some values of amplitude (ω), when the frequency of encounter with bacteria (Φ) is intermediate (supplemental figure 6). The exception occurred when induction was optimized with Φ = 0.1. In this case, the induced defense performs poorly when compared to a constitutive defense for most frequencies and amplitudes of exposure. A limitation of this analysis is that fly-bacteria encounters are predictable when there is a deterministic sinusoidal input of bacteria, which is not biologically realistic.

To model unpredictable encounters, we compared the fitness of induced and constitutive defenses upon stochastic encounters with bacteria. To those ends, we simulated stochastic fly-bacteria interactions using random walks of a fly in a 100 × 100 grid with differing densities and patchiness of bacteria (Figure 2). For each combination of bacterial density and patchiness values, we report the average relative fitness of constitutive versus induced defenses across 1,000 random walks for each value of Φ and *k*_0_ used to optimize the induced defense.

First, we tested a constitutive defense against an induced response with equal degradation rates (λ_2_) for all proteins involved in the Imd pathway (i.e., an 11-parameter model). For most values of Φ, induced defenses had higher fitness values than constitutive defenses in environments with a heterogeneous distribution and low-to-intermediate density of bacteria (Figure 3). For the highest bacterial proliferation rate (*k*_0_ = 0.5) and Φ < 0.1, induction had the highest relative fitness when bacterial density was lowest and distributed with maximal heterogeneity (Figure 3). For other bacterial proliferation rates (*k*_0_ = 0.1 and *k*_0_ = 0.2) and Φ < 0.1, the relative fitness for induction was highest in environments with an intermediate density of bacteria and a heterogeneous distribution (Figure 3). The only exception to these patterns was when the induced response was optimized with Φ = 0.1 and tested against bacteria with an intermediate (*k*_0_ = 0.2) or a high (*k*_0_ = 0.5) proliferation rate (Figure 3). For these parameter combinations, induction performed best at higher bacterial densities. The poor performance of the induced defense optimized with Φ = 0.1 in the stochastic model is consistent with results from the sinusoidal model of fly-bacteria encounters (supplemental figure 6). In general, induction almost always performed best, relative to constitutive defense, in environments with heterogeneous bacterial distributions. However, the effect of bacterial density on the relative fitness of induced and constitutive defenses depended on the values of Φ and *k*_0_ used to optimize the induced defense.

**Figure 3.**
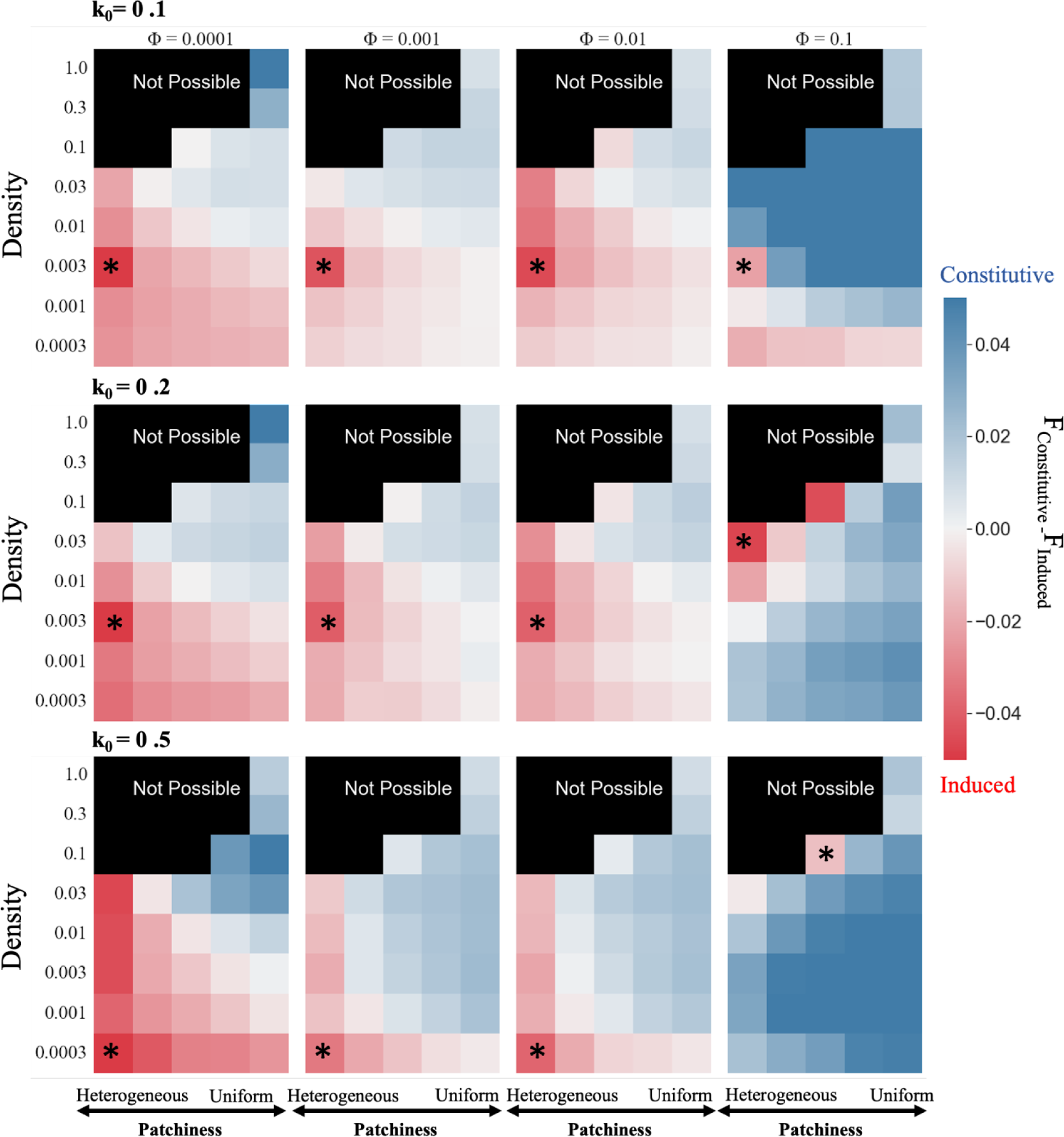
Induced responses tend to outperform constitutive defenses when flies inhabit environments with low densities and heterogeneous distributions of bacteria. The relative fitness of constitutive versus induced strategies are shown as heatmaps for environments with different densities (*d*) and patchiness (*p*) of bacteria. Patchiness values (*p*) for each cell are reported in Supplemental Table 2. Red cells indicate that the fitness of the induced response (F_Induced_) is higher than fitness of the constitutive defense (F_Constitutive_), and blue cells indicate that F_Constitutive_ > F_Induced_. Heatmaps in the same column show induced responses that were optimized with the same frequency of the sinusoidal input of bacteria (Φ), and heatmaps in each row show the results for the same proliferation rate of bacteria (*k*_0_). Asterisks show the environment in which the induced defense has the highest relative fitness.

Next, we evaluated how the fitness function affects the relative performance of the induced and constitutive strategies. To this end, we optimized the induced defense by assuming that the production of proteins involved in Imd signaling is not costly, and the only cost comes from bacterial proliferation and production of AMPs (Eq. 11; supplemental figure 7). This assumption increases the relative fitness of the induced defense, such that induction generally outperforms constitutive defense regardless of the bacterial environment (supplemental figure 8A). Consistent with our results using a more costly fitness function (Figure 3), however, induced defense has the highest relative fitness in environments with intermediate densities and heterogeneous distributions of bacteria (supplemental figure 8A). Also, the relative fitness of induction is higher in heterogeneous and low density environments, and induction performs worst in high density and uniform environments, regardless of the fitness function (Figure 3; supplemental figure 8A). As before, induction performs worst when optimized with Φ = 0.1 (supplemental figure 8B). Because we observe the same overall pattern of relative fitness of induced and constitutive strategies using both fitness functions, we conclude that our results are robust to the choice of the fitness function. For the remainder of the manuscript, we use the fitness function in which Imd signaling is costly (Eq. 10).

The analysis above assumes identical degradation rates (λ_2_) for Imd signaling pathway proteins (it is an 11-parameter model), and we tested the effects of this assumption. To that end, we optimized a model of induction with different parameters for degradation rates (16-parameter model). We used two approaches to optimize the 16-parameter model, both of which start from the parameter values optimized in the 11-parameter model. In the first approach, we search for the optimum fitness by allowing small changes for all 16 parameters. Using this approach we found that the six degradation rate parameters converge to the single degradation parameter in the 11-parameter model (λ_2_) (supplemental figure 9). This suggests that there is no benefit to separately optimizing each degradation rate. In the second approach, we found the optimum induced defense by varying the six degradation parameters, while fixing the other 10 parameters. Using this approach we found optimized induced defenses with different degradation rates for proteins involved in Imd signaling (supplemental figure 9). We found that the induced response has the highest performance when compared to constitutive defense under the same conditions for both the 11-parameter and 16-parameter models (supplemental figure 10). We therefore conclude that the 11-parameter model is sufficiently complex to capture the costs and benefits of an induced immune response, and our subsequent analyses focus on the 11-parameter model.

We further examined the effects of bacterial density, patchiness, and proliferation rate on the performance of the induced and constitutive defenses. To this end, we calculated the proportion of environments with different combinations of density and patchiness in which induction outperforms constitutive defense for a given bacterial proliferation rate and optimization of the induced response. This was done by calculating the fraction of red cells out of the total number of cells for each heatmap in Figure 3 for the stochastic model, and supplementary Figure 6 for the sinusoidal model, which we refer to as the “proportion of induced wins” (PIW). Under the stochastic models, induction performed best (i.e., highest PIW) when bacterial proliferation rates were low or medium (Figure 4A–B). As bacterial proliferation rates increased, constitutive defense performed better in most environments (i.e., PIW decreased). The exception to this rule was when Φ = 0.0001, in which case PIW remained fairly constant regardless of the bacterial proliferation rate.

**Figure 4.**
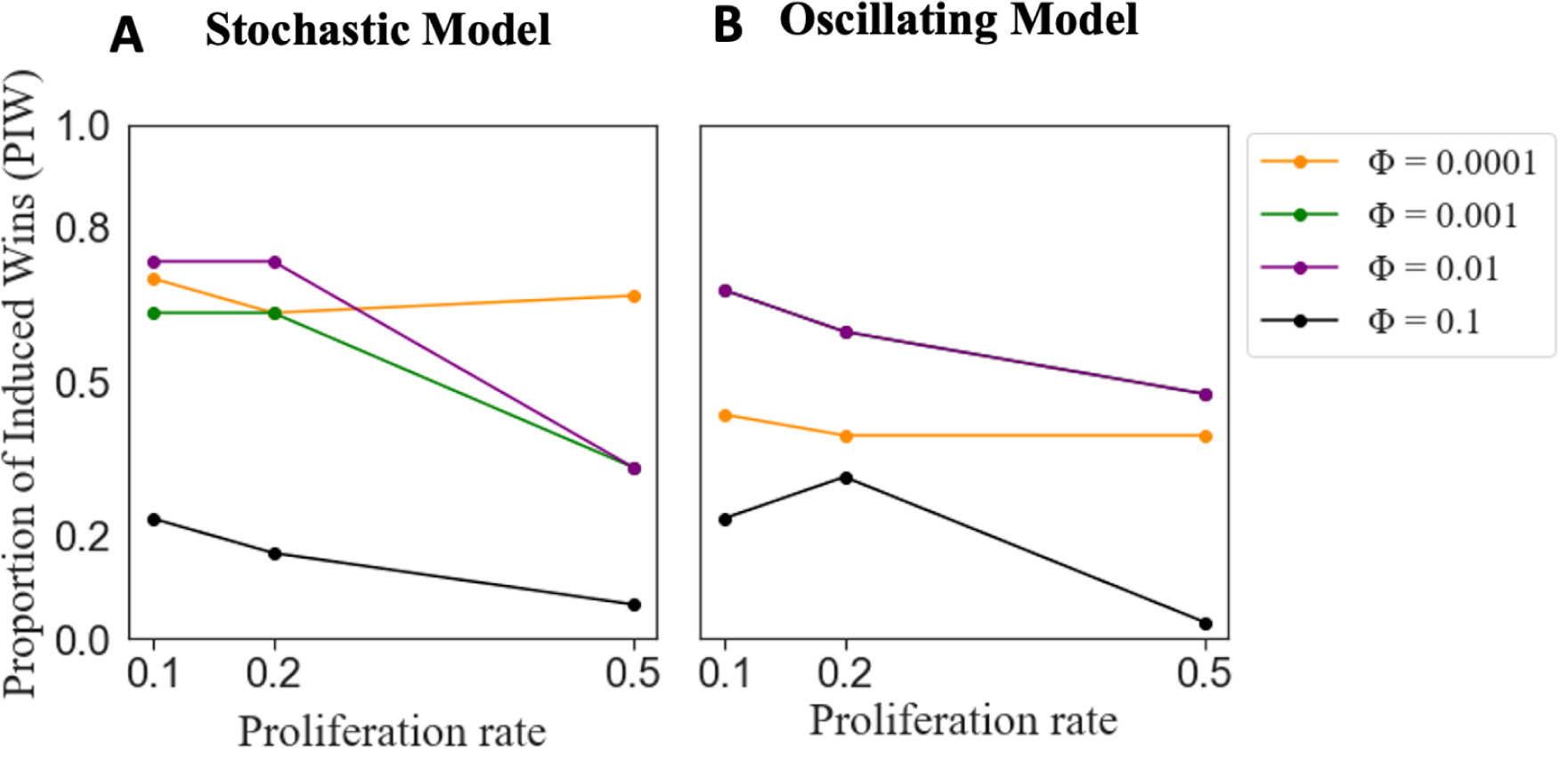
Proportion of induced wins (PIW) (Y-axis) for different proliferation rates (*k*_0_) of bacteria (X-axis) under the stochastic (A) and sinusoidal oscillating models (B). PIW measures the proportion of environments in which an induced response outperforms constitutive defense. The results are shown for induced response optimized in environments with varying frequencies of encounter with bacteria (Φ). The green and purple lines (Φ = 0.01 and Φ = 0.001) on panel B overlap.

Results under the sinusoidal oscillating model were consistent with the stochastic models in three ways. First, the performance of the induced defense optimized with Φ = 0.0001 was relatively unaffected by the bacterial proliferation rate under all models. Second, under both stochastic and oscillating models, when optimized with Φ = 0.1, induction performs poorly against a constitutive defense, especially when the bacterial proliferation rate is high. Finally, under both models, for intermediate values of Φ (Φ = 0.001 and Φ = 0.01), higher proliferation rates reduced PIW (Figure 4).

Our analysis is unlikely to be biased by the parameters we selected in Figures 3 and 4. We explored specific values for bacterial density and patchiness in the stochastic model; amplitude and frequency in the sinusoidal model; and proliferation rate in both models. If we had sampled more environments with a high bacterial density, PIW would be lower. However, the observed trends in Figure 4 are unaffected by our parameter values because we are comparing the relative performance of induced and constitutive defenses across different values of Φ and *k*_0_using fixed values of density and patchiness. In other words, the relative performance (i.e., change in PIW) across proliferation rates (Figure 4) should be robust to the parameters selected in the heat maps (Figure 3). Because our results are consistent under both stochastic and sinusoidal models, we will proceed only with the stochastic model because it allows us to consider fitness differences between the induced and constitutive defenses when fly-bacteria encounters are unpredictable.

### Inhabiting multiple types of environments favors induction

We next evaluated the performance of induced and constitutive defenses when a fly inhabits multiple environments. We were specifically interested in whether experiencing multiple types of environments (i.e., different bacterial densities and patchiness) favors induction, even when each individual environment favors constitutive defense. For that reason, we only sampled from environments in which constitutive defense outperforms induction (blue cells in Figure 3), and then we measured the relative fitness of induced and constitutive defenses when flies inhabit multiple environments that were each sampled with equal probabilities. We calculated PIW when flies experience multiple bacterial environments for induced strategies that were optimized with different frequencies of bacterial input (Φ), across different numbers of possible environments, and at three different bacterial proliferation rates (*k*_0_ = 0.1, 0.2, 0.5).

We found that inhabiting multiple environments favored induction over a constitutive defense. PIW generally increased as the number of environments increased, often reaching its maximum value of 1. Whether PIW reaches its maximum depends on the Φ used to optimize the induced defense. PIW reached one for induced defenses optimized with lower frequencies of bacterial input (Φ = 0.0001, Φ =0.001, and Φ = 0.01), regardless of the bacterial proliferation rate (Figure 5, first three rows). In addition, the rate of increase of PIW was greater for the lower proliferation rates (*k*_0_ = 0.1 or 0.2) than the higher proliferation rate (*k*_0_ = 0.5). PIW > 0.5 indicates that induction is favored because induced defenses win more than they lose. In general, PIW>0.5 when the number of environments exceeded either 2 or 3, for most combinations of proliferation rates (*k*_0_) and Φ values (Figure 5). In contrast, when induction was optimized with a high frequency of bacterial input (Φ = 0.1), PIW remained low for both low and high bacterial proliferation rates (*k*_0_ = 0.1 or 0.5).

**Figure 5.**
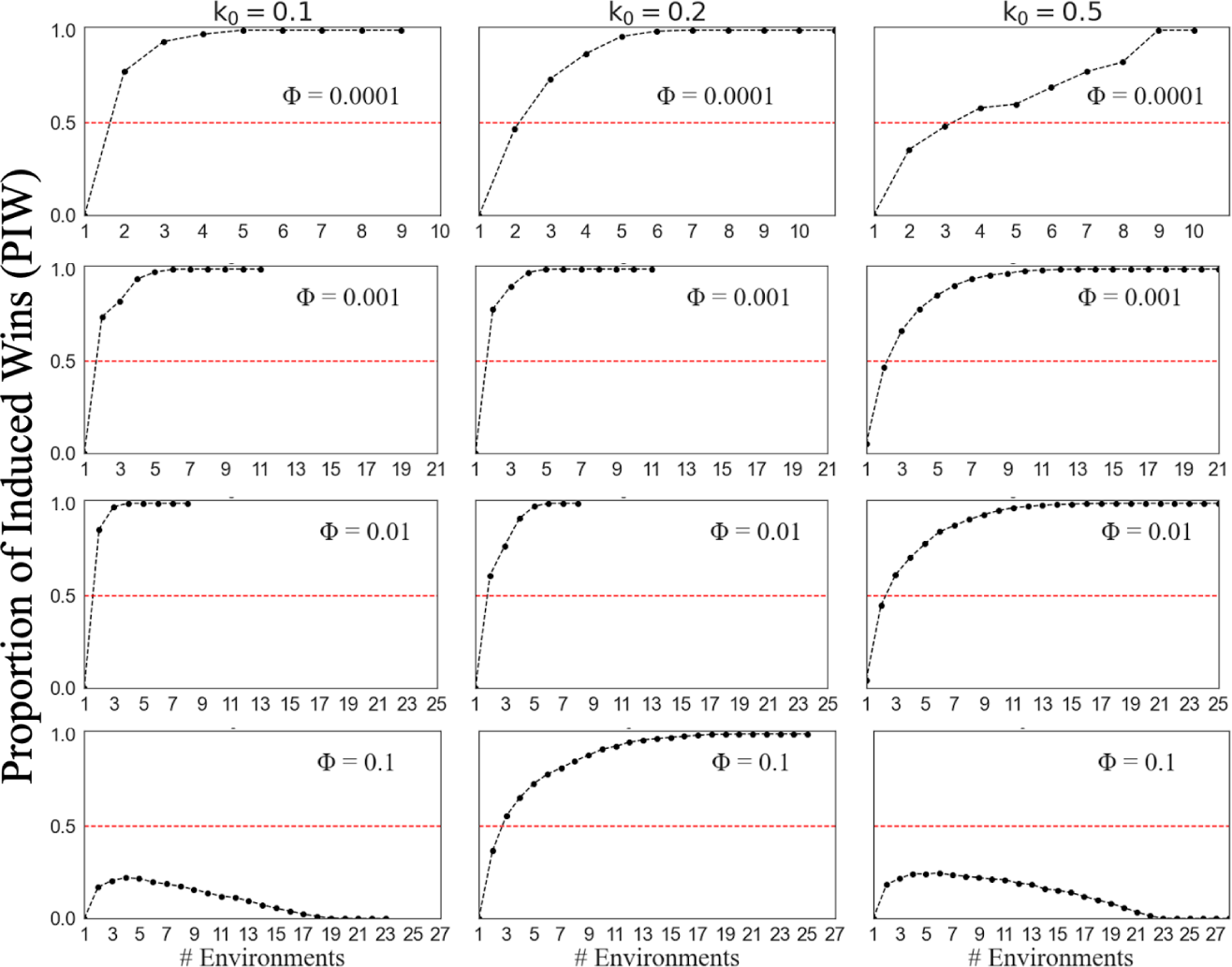
Induction outperforms constitutive defense as the number of different environments a fly inhabits increases. The proportion of times induction outperforms constitutive defense, PIW (Y-axis), is plotted against the number of possible environments that are sampled (without replacement). Graphs in a column have the same bacterial proliferation rate (*k*_0_), and graphs in a row have the same Φ value used to optimize the induced model. Induction outperforms constitutive defense above the dashed line (PIW>0.5), and constitutive defense performs better below the dashed line (PIW<0.5).

Next, we explored how the frequency of time the fly spends in two different environments affects the relative fitness of induced and constitutive defenses. As above, we calculated PIW when we sampled from individual environments where constitutive defense has a higher fitness than induction. We found that PIW was greatest when the two environments occur at equal frequencies (*q* = 0.5), regardless of the frequency of the bacterial input used to optimize the induced response (supplemental figure 11). PIW decreased to 0 as the frequency of time spent in one environment increased.

### Negative regulators of Imd signaling affect the fitness of induction depending on the bacterial proliferation rate

We compared the concentrations of negative regulators of the Imd signaling pathway in response to bacteria with different proliferation rates (*k*_0_) for induced defenses optimized with different frequency of exposure to bacteria (Φ). This is motivated by the observation that the fitness of induced defenses that are optimized with different values of Φ depends on the bacterial proliferation rate. We compared the induced defenses optimized with either Φ = 0.0001 or Φ = 0.01. This is because induction optimized with Φ = 0.0001 offers the best defense against bacteria with a high proliferation rate (Figure 4). In contrast, optimization with Φ = 0.01 offers the best response against bacteria with a low proliferation rate (Figure 4). We aimed to understand how the optimization of these different induced defenses shapes the production of negative regulators, thereby affecting their performance across bacterial environments.

We found that for low bacterial proliferation (*k*_0_ = 0.1), induction optimized with Φ = 0.01 (which is the best optimization for *k*_0_ = 0.1) produces more negative regulators that reduce the input to the Imd pathway (PGRP-LB and Pirk) compared to an induced defense optimized with Φ = 0.0001 (Figure 6). Our results are consistent between heterogeneous and uniform environments (supplemental figure 12). Therefore, we hypothesized that investing in negative regulators that reduce the input to the signaling pathway is beneficial against bacteria with low proliferation rates. We tested this hypothesis by measuring the fitness of induced defenses upon changing the parameter values that determine the production rates of PGRP-LB and Pirk. To this end, we first chose the induced defense with the worst performance, which is optimized by Φ = 0.1 and *k*_0_ = 0.5 (Figure 4). Next, we tried to improve the fitness of this induced defense by multiplying the two parameter values that determine the production rates of PGRP-LB and Pirk (β_3_ and β_4_ in Equations 6 and 7) by a constant (γ). We measured improvement by calculating PIW upon changing the two parameter values (Figure 7). We found that for a low bacterial proliferation rate, the maximum PIW is achieved for larger rates of production of PGRP-LB and Pirk (γ = 0.33) compared to the high bacterial proliferation rate (γ = 0.25) (Figure 7). This is consistent with the hypothesis that the induced defense with a higher rate of production of PGRP-LB and Pirk is more successful against bacteria with a low proliferation rate.

**Figure 6.**
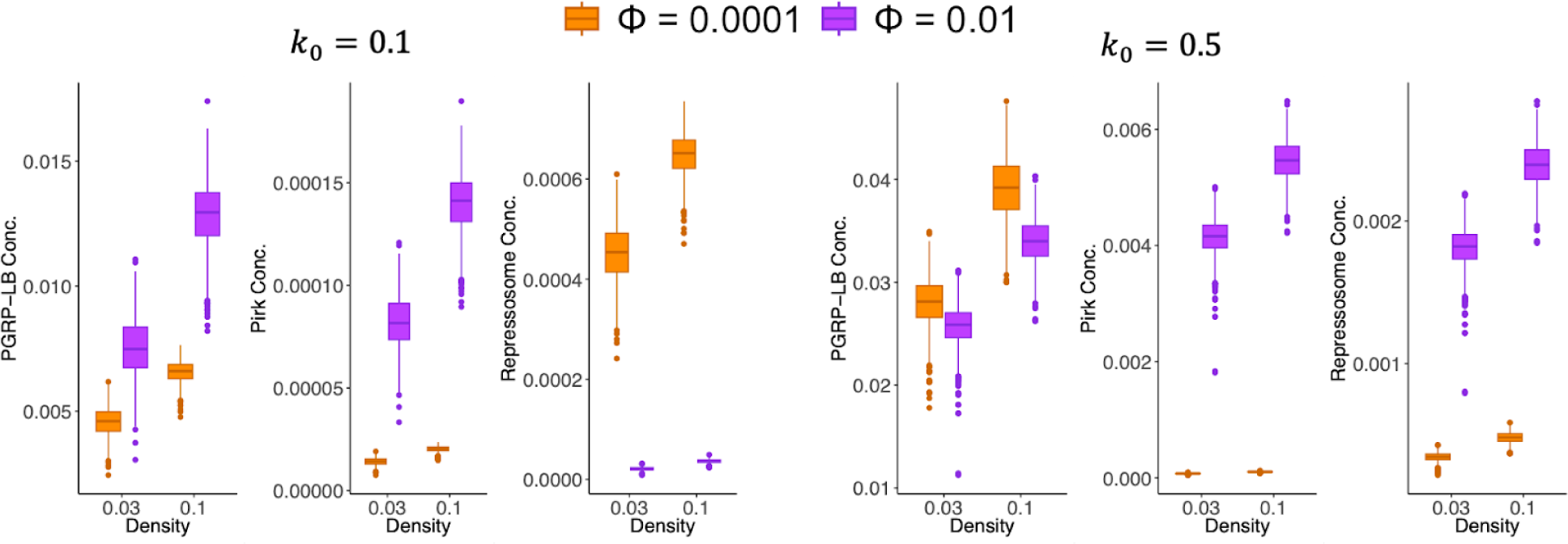
The box plots show the concentration of proteins involved in negative regulation of the Imd signaling pathway (Y-axis) in environments with two different densities (d) of bacteria (X-axis) that posses different proliferation rates (*k*_0_) across 1,000 simulations. Induced defenses were optimized with different frequencies of bacterial encounters (Φ). The distribution of bacteria in all analyses is heterogeneous (*p* = 1).

**Figure 7.**
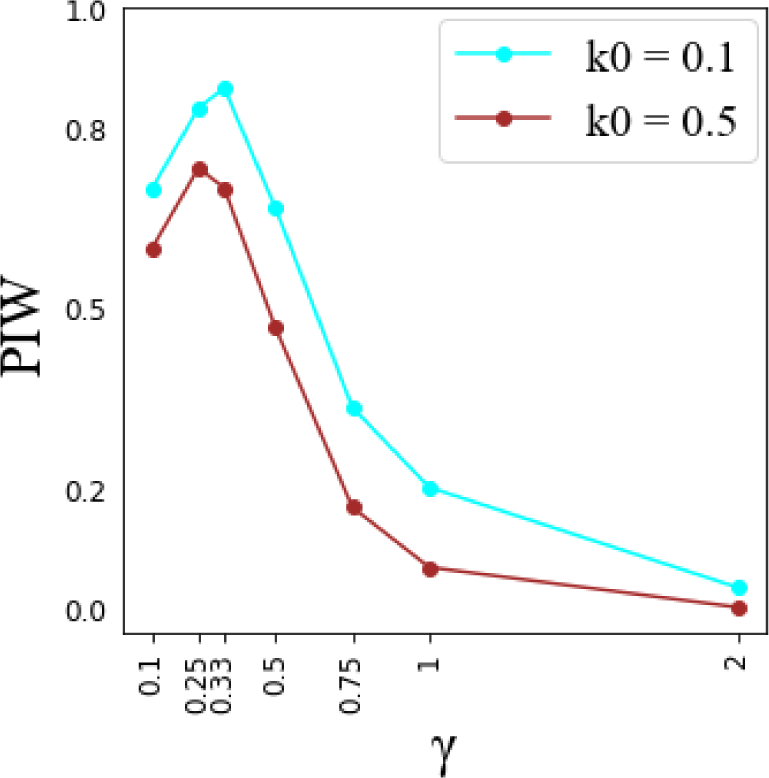
Calculation of proportion of induced wins (PIW) for different parameter values that control the production rates of PGRP-LB and Pirk. The parameter values are changed by multiplying the optimized values for Φ = 0.1 by a constant (γ) (X-axis). The analysis is done for low (cyan line) and high (brown line) bacterial proliferation rates (k0).

Next, we sought to understand what constitutes an effective immune defense against bacteria with a high proliferation rate. Because we found that higher rates of PGRP-LB and Pirk production were detrimental against bacteria with a high proliferation rate (Figure 7), we focused our attention on the third negative regulator, namely, the repressosome complex. We found that for the high bacterial proliferation rate (*k*_0_ = 0.5) the induced defense optimized with Φ = 0.0001 (which is the best optimization for *k*_0_ = 0.5) produces less repressosome than the induction optimized with Φ = 0.01 (Figure 6). However, the binding energy for repressosome was lower (a more effective binding) for induction optimized with Φ = 0.0001 (*Z_S_* = 5 as opposed to *Z_S_* = 5.1 for Φ = 0.01). In our model, the fitness of an induced defense is influenced by both the production rate of the repressosome and its binding energy. Therefore, we needed to look beyond the concentration of repressosome to measure its effect on fitness. To this end, we try to improve the poor immune response optimized with Φ = 0.1 and *k*_0_ = 0.5 by multiplying the binding energy of the repressosome complex (*Z_S_* in Eq. 9) by a constant (δ). We did not change the rate of repressosome production because by changing the binding energy we can examine the effect of repressosome on fitness without confounding it with the cost of the production of the repressosome itself. We found that the PIW is consistently reduced for larger binding energy values (poor binding) regardless of the bacteria proliferation rate (supplemental figure 13). Hence, assessing PIW does not provide insight into the potential benefits of repressosome upon induction. We next measured the fitness of induced defenses in individual environments with a unique combination of bacterial density and patchiness upon changing the binding energy of the repressosome complex. We found that the induced defense reaches its maximum fitness across more environments for lower binding energy values (e.g., δ = 2 or 4) when the bacterial proliferation rate is low (*k*_0_ = 0.1) than when it’s high (*k*_0_ = 0.5) (Figure 8A). However, when we reduced the rate of PGRP-LB and Pirk production (γ = 0.5), we observed the opposite pattern (Figure 8B). For lower production rates of PGRP-LB and Pirk, the induced defense reaches its maximum fitness across more environments for lower binding energy values (e.g., δ = 1) when the bacterial proliferation rate is high (*k*_0_ = 0.5) than when it’s low (*k*_0_ = 0.1) (Figure 8B). Thus our results suggest that damping the immune response using repressosome is effective against bacteria with a high proliferation rate if PGRP-LB and Pirk are produced at low levels (Figure 8B).

**Figure 8.**
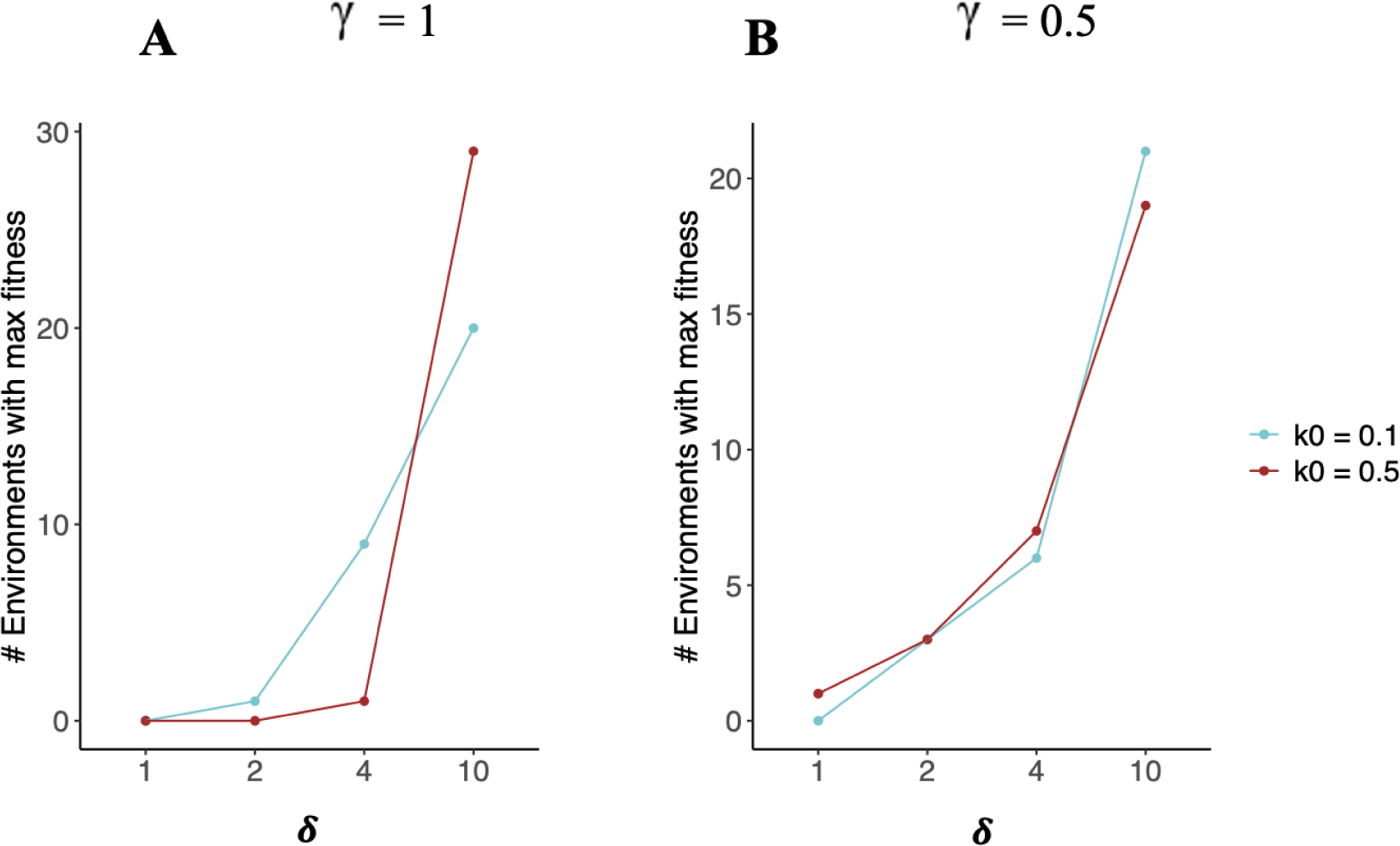
Number of environments (out of a total of 30) in which induced defense reaches its maximum fitness (Y-axis) for four repressosome binding energy values (X-axis). The repressosome binding energy is manipulated by multiplying by a constant (δ) for an induced defense that is optimized with Φ = 0.1 and *k*_0_= 0.5. The analysis is done for low (cyan line) and high (brown line) bacterial proliferation rates (k0). (A) High production of PGRP-LB and Pirk (γ = 1). (B) Low production of PGRP-LB and Pirk (γ = 0.5).

## Discussion

We modeled constitutive and induced immune defenses of *D. melanogaster* (Figure 1), and we compared their relative fitness in environments with different densities and patchiness of bacteria (Figure 2). Our model predicts that induction is favored in environments where bacterial exposure is uncertain, which can be caused by heterogeneous distributions of bacteria (Figure 3) or fluctuations in the bacterial density and patchiness (Figure 5). However, the specific benefits of induced defense depend on the proliferation and distribution of bacteria used to optimize the model and to which the flies are exposed (Figure 4). Interestingly, we found that induction is favored when flies experience multiple bacterial environments, even when those individual environments favor a constitutive defense in isolation (Figure 5, supplemental figure 11). Our results are robust under most approaches for optimizing the parameters of the induced response (Figure 3, supplemental figures 8A and 10), and support the hypothesis that uncertainty over encountering bacteria favors an induced immune response (Adler & Karban, 1994; Hamilton et al., 2008). By including the mechanistic details of three specific negative regulators of induction, we showed that investing in negative regulators that reduce the input to the system results in an effective defense against bacteria with low proliferation rates (Figure 7). On the other hand, our model predicts that investing in negative regulators that reduce the output (AMP) produces effective defense against bacteria with a high proliferation rate (Figure 8). Thus, our results shed light on the evolution of negative feedback mechanisms that control immune responses by reducing either input to the signaling pathway or the output.

### Uncertainty favors an induced immune response

Uncertainty of bacterial exposure can come from multiple sources, and we find that the interactions of these different sources can affect the extent to which different types of uncertainty favor induction. For example, the distribution of bacteria in the environment can create uncertainty, which favors induction in specific combinations of bacterial densities and patchiness. The uncertainty of exposure is maximal when bacteria are at intermediate densities with heterogeneous distributions because encounters are frequent yet unpredictable (Figure 5). We found that induction is most favored at this maximal uncertainty (Asteriks in figure 3, supplemental figure 3B, 8A, and 10). Other combinations of density and patchiness can result in certainty of exposure (Figure 9). For example, certainty of encountering bacteria occurs in environments with a high density and uniform distribution of bacteria, which ensures frequent encounters with bacteria. We found that environments with high bacterial densities and uniform distributions do indeed favor a constitutive defense (top right of heat maps in Figure 3, supplemental figure 3B, 8A, and 10). Uncertainty in our model can also come from inhabiting multiple environments with different combinations of bacterial densities and patchiness. A key finding of our model is that induction is favored when flies experience multiple environments, even when a constitutive defense is favored in each individual environment (Figures 5 and supplemental figure 11). In particular, a high level of uncertainty for choosing between 2 environments, i.e., equal probabilities of each environment, is especially favorable for induction (supplemental figure 11). Even though it is well established that uncertainty favors induction, our model is unique in predicting that induction can be favored upon environmental fluctuations even if constitutive defense outperforms induced defense in each individual environment.

**Figure 9.**
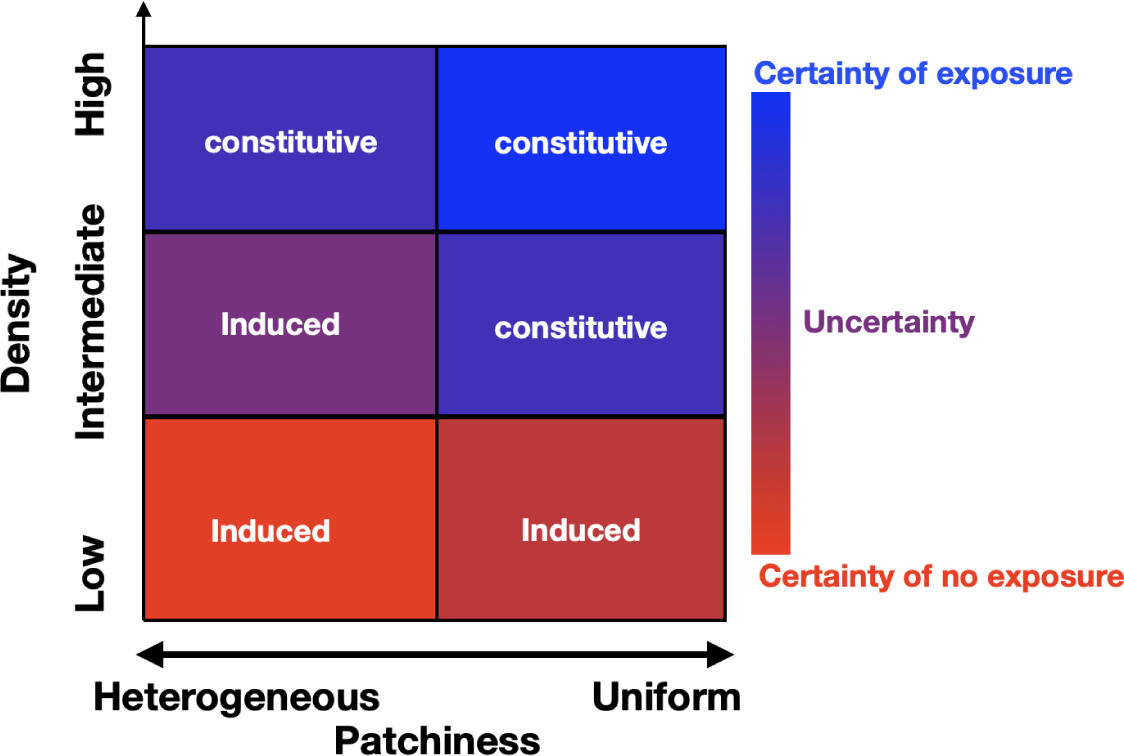
Certainty and uncertainty of encountering bacteria in environments with different density and patchiness of bacterial populations. Certainty of exposure is shown with shades of blue, and certainty of no exposure is shown with shades of red. Environments that favor constitutive defenses are specified by “constitutive”, and environments that favor induced defenses by “induced”.

There are important parallels between the uncertainty that arises from multiple possible environments in our model and selection pressures that act on AMP protein sequences. Balancing selection can maintain sequence variation in *Drosophila* AMP genes (Unckless & Lazzaro, 2016). Balancing selection on AMPs likely occurs as a result of fluctuation of bacterial populations in environments inhabited by flies (Abdul-Rahman et al., 2021). Such fluctuating environments are similar to those that favor induced defenses in our model. Therefore, uncertainty might have been responsible for both the prevalence of induced expression of AMPs and AMP sequence variation in *Drosophila*.

We identified one important exception to the rule that certainty favors a constitutive defense. When bacterial density is low and distribution is heterogeneous, fly-bacteria encounters are rare, which creates high certainty of no exposure (Figure 9). In contrast to certainty of bacterial exposure (which favors constitutive defense), certainty of no exposure favors an induced defense. This can be seen in environments with low bacterial density and heterogeneous distributions, which favor induction because of a low frequency of fly-bacteria encounters (bottom left of heat maps in Figure 3, supplemental figure 3B, 8A, and 10). We interpret this to mean that, when there is high certainty of no exposure, a constitutive defense is costly without much benefit, which favors induction.

### Bacterial proliferation rate affects the benefits of induction and negative regulators

We also find that bacterial proliferation rate affects the benefit of an induced response in uncertain environments. Hamilton *et al*. (2008) showed that induction is favored when the bacterial proliferation rate is unknown. Other work showed that a higher pathogen growth rate favors constitutive defense over induction (Kamiya et al., 2016; Shudo & Iwasa, 2001). However, previous work ignored structural differences in bacterial distributions (here modeled as patchiness and density) when calculating effects of bacterial proliferation on the relative performance of the induced and constitutive defenses. One of the benefits of our model is that it allows for separate and combined study of the effect of bacterial proliferation rate and uncertainty on the relative performance of induced and constitutive defenses.

Our results show that the effect of bacterial proliferation rate on the relative fitness of induced and constitutive defenses depends on the bacterial density and distribution. For example, we found that the relative fitness of the induced defense is highest against bacteria with low proliferation rates if bacterial density is intermediate and distribution is heterogeneous (Figure 3). However, we found that a higher bacterial proliferation rate strongly selects for an induced defense if the bacterial density is low and the distribution is heterogeneous (Figure 3). The latter result makes intuitive sense because, in environments with low bacterial densities and heterogeneous distributions, fly-bacteria interactions are rare, and an immune response is not needed at most times. However, if the bacterial proliferation rate is high, upon ingestion of bacteria, a high level of AMP production is needed to eliminate the pathogen. In this situation, low constitutive AMP expression is not effective because it cannot efficiently eliminate the bacteria. High constitutive expression of AMPs, on the other hand, can eradicate a bacterial infection, but the cost of constitutive production of AMPs outweighs the benefits because AMPs are not needed at all times. An induced defense, however, which produces large amounts of AMPs upon demand, is optimal for rare encounters with bacteria that have a high proliferation rate.

Our strategy for optimizing parameter values against different frequencies of bacterial inputs (Φ) is also informative of the relative fitness of induced and constitutive defenses across bacterial proliferation rates. Our results agree with previous findings which demonstrated that the evolution of induction is more likely when the frequency of host-pathogen interactions is low (Hamilton et al., 2008). Therefore, we would expect induction to be more effective if it is optimized in environments where host-pathogen interactions are not frequent. Consistent with this prediction, we found that the relative fitness for the induced defense is lower when optimized with a high frequency of bacterial input (i.e., Φ = 0.1), especially for higher bacterial proliferation rates (Figure 4).

Our model also predicts that bacterial proliferation rate affects the benefits of producing different negative regulators of the Imd pathway. Specifically, we found that production of negative regulators that reduce the input into the Imd signaling pathway (PGRP-LB and Pirk) increases the relative performance of an induced response against bacteria with low proliferation rates (Figure 7). On the other hand, production of more repressosome complexes—which compete with Relish for binding to the promoter of AMP genes—results in an effective immune response to bacteria with a high proliferation rate (Figure 8) (Relish is the transcription factor that induces AMP expression in the Imd pathway). We hypothesize that investing in negative regulators that reduce the input to the system can rapidly shut down the pathway and prevent wasteful investment in immunity. This is an effective strategy against bacteria with low proliferation rates because it imposes less fitness costs on the host. On the other hand, when defending against bacteria with a high proliferation rate, investing in negative regulators that reduce the output (AMPs) is more effective because it lowers the cost of AMP production without entirely shutting down the pathway. Unlike PGRP-LB and Pirk, the repressosome does not reduce the concentration of active Relish. In this way, active Relish is available for a timely response upon subsequent encounters with rapidly dividing pathogens. Previous theoretical work that did not simultaneously model the multiple negative feedback loops of the Imd signaling pathway and bacterial proliferation rates could not have predicted these results. For example, Ellner et al. (2021), who provided a detailed model of the Imd pathway, did not explicitly model the action of three negative feedback loops (PGRP-LB, Pirk, and the repressosome complex), and therefore could not observe their different effects. Future empirical studies could test how each negative regulator of the Imd signaling pathway affects the survival of flies when they encounter bacteria with different proliferation rates.

The finding that negative feedback mechanisms operating at different steps of the signaling pathway are beneficial against bacteria with different proliferation suggests that uncertainty not only favors induction but also shapes the mechanisms that regulate induction. In the natural environment, flies encounter an array of bacteria that multiply at varying rates (Duneau et al., 2017). Thus, the evolution of immune responses is not only influenced by the unpredictability of bacterial encounters but also by uncertainty with regard to rates of bacterial proliferation. This might explain the fact that immune responses are often regulated at different steps in the signaling pathway. For example, interferon signaling is regulated by downregulation of cell surface receptors and inhibition of downstream transcription factors (Porritt & Hertzog, 2015). Our study contributes to understanding the prevalence of negative regulators with distinct functions in immune signaling pathways. In addition, given the widespread prevalence of negative feedback loops beyond immune signaling pathways, the insights derived from this model could potentially extend to other signaling pathways as well (El-Samad, 2021; Perrimon & McMahon, 1999; Shilo, 2014; Yu et al., 2008).

---

**Supplemental figure 1.**
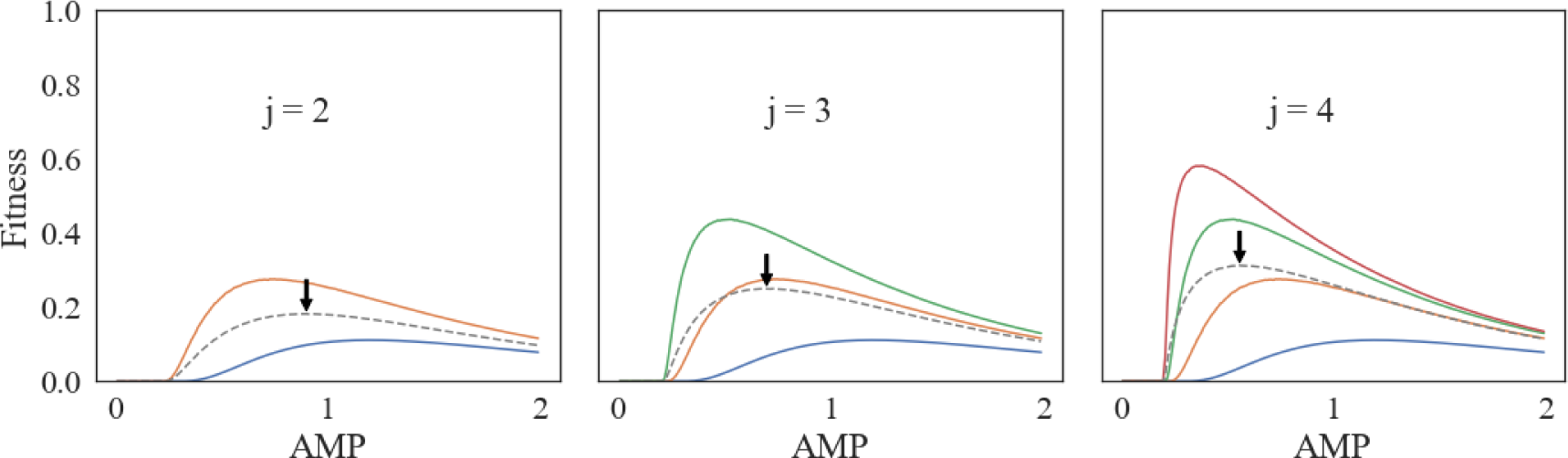
Finding the best constitutive defense when a fly inhabits *j* environments, with different density and patchiness of bacteria, with equal probability. Fitness is shown on the Y-axis in environments with different density and/or patchiness of bacteria (different colors) for varying levels of constitutive AMP expression (X-axis). The dashed line represents the fitness if the fly randomly encounters *j* environments each with a probability 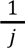. The arrow shows the optimum fitness upon random encounter with *j* environments.

**Supplemental figure 2.**
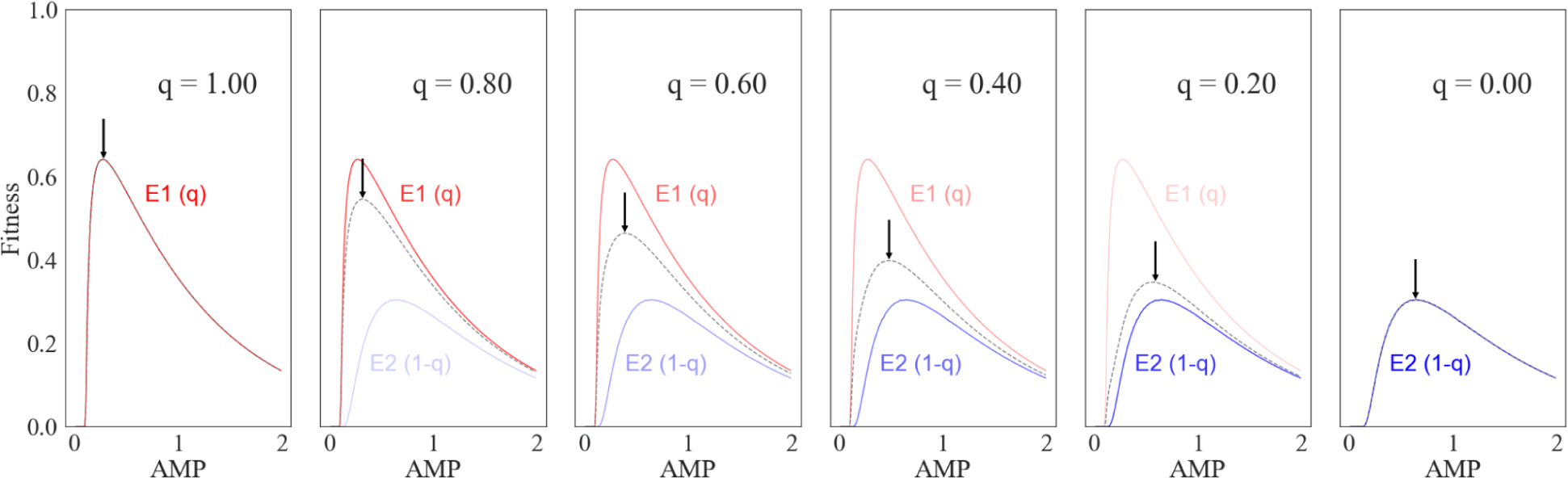
Optimum fitness of constitutive defense (shown with an arrow) when flies inhabit two environments, E1 and E2, with probability *q* or 1 − *q*, respectively. Red lines represent the fitness in E1 and blue lines represent the fitness in E2. The dashed line is the fitness when the fly inhabits both environments with different probabilities. The optimum fitness is shown with an arrow.

**Supplemental figure 3.**
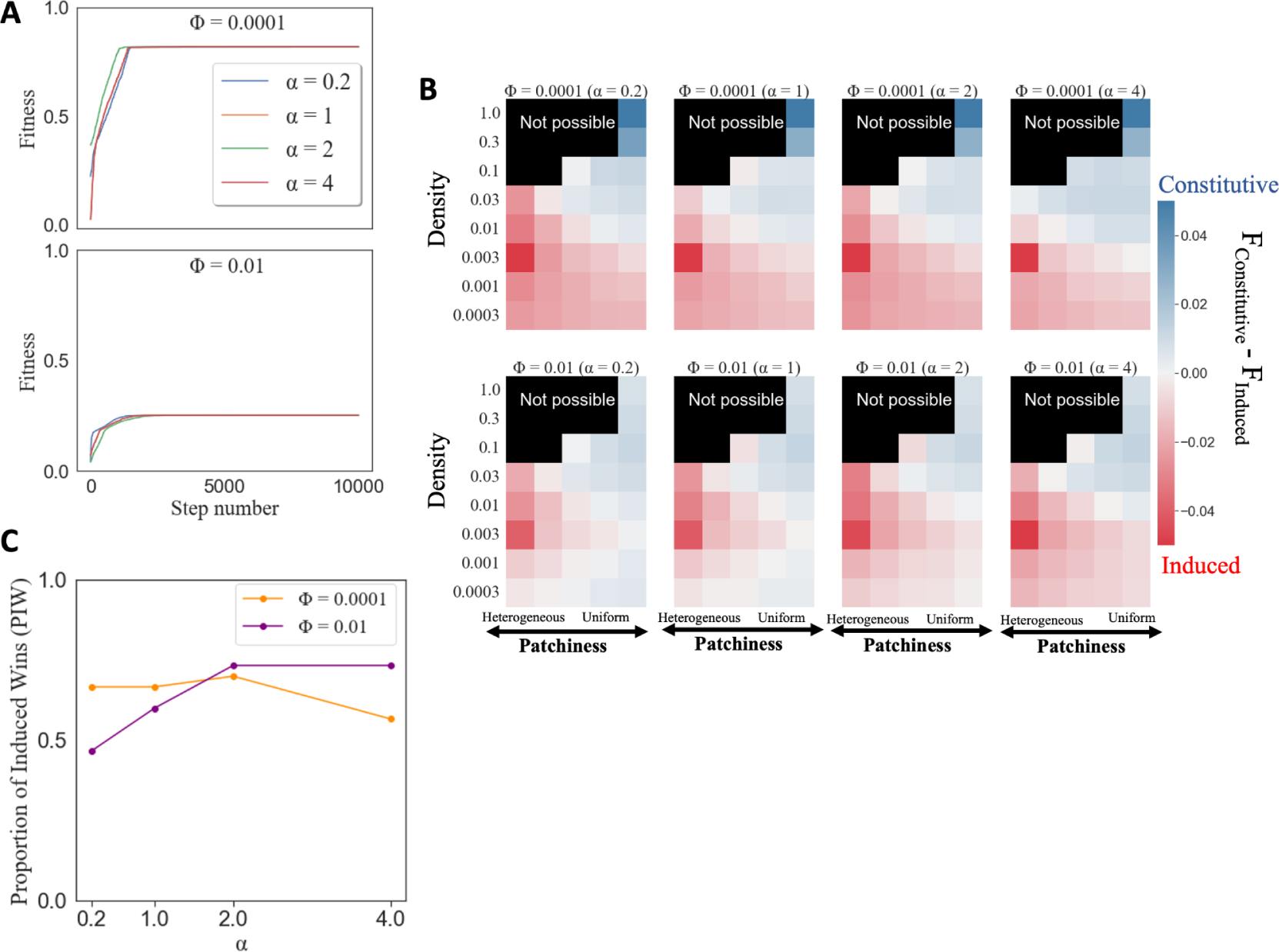
Results for optimization of induced defense for four values of α (0.2, 1, 2, and 4). **A:** The fitness value (Y-axis) at every step of optimization (X-axis) for optimization that resulted in the highest fitness amongst 2048 simulations. Optimizing induced defense for different values of α does not change the optimum fitness value for two frequencies of input to the system (Φ = 0.0001 and Φ = 0.01). **B:** The relative fitness of induction vs constitutive defense (color bar) in environments with different patchiness (X-axis) and bacterial density (Y-axis). Induction has a higher fitness in red-colored cells than the best constitutive defense. The highest relative fitness is shown with an asterisk. Induction is preferred in heterogeneous environments with low bacterial density regardless of the value of α. **C:** Comparing the proportion of induced wins (Y-axis) across different values of α (X-axis) for two values of Φ. The proportion of induced wins is highest for α = 2 regardless of the value of Φ.

**Supplemental figure 4.**
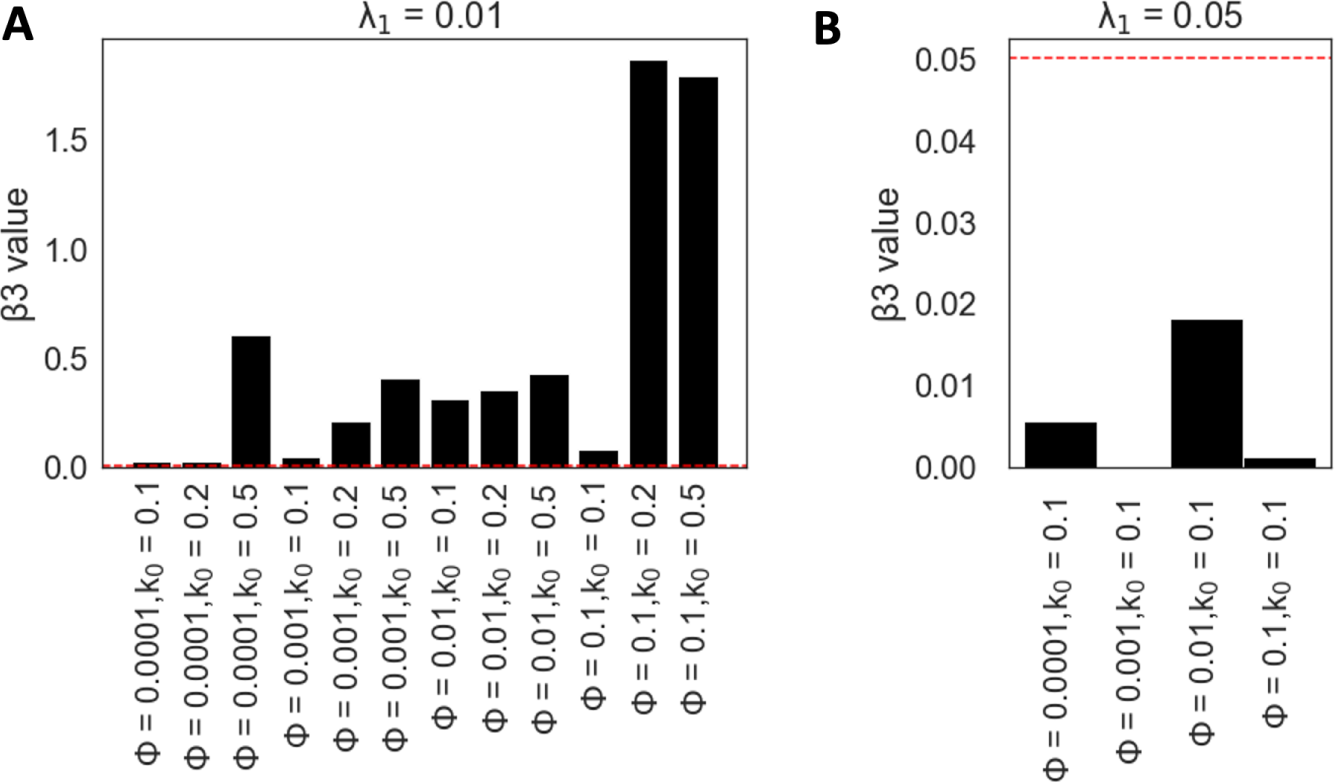
Optimization of the induced defense with two values of λ_1_. The X-axis are the parameters affecting the production of PGRP-LB (β_3_), and the Y-axis is the value of β_3_in the optimized model. **A:** Optimization of the induced defense with λ_1_= 0.01 using different values of Φ and *k*_0_. The red dashed line is the value of λ_1_(0.01). **B:** Optimization of the induced defense with λ_1_= 0.05 using different values of Φ. The red dashed line is the value of λ_1_(0.05).

**Supplemental figure 5.**
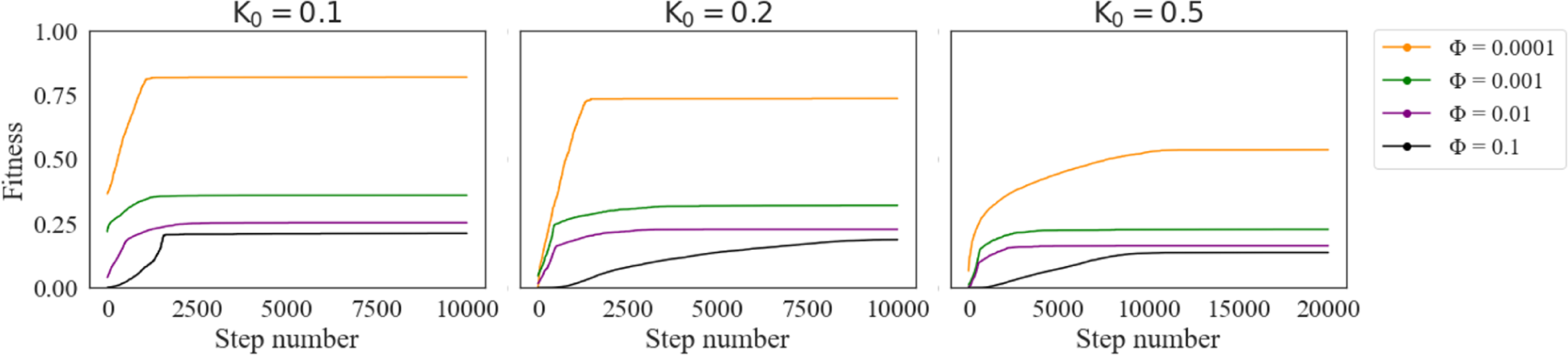
Optimization of the induced response using an oscillating deterministic input (sinusoidal) with different frequencies (Φ) for three different proliferation rates of bacteria inside the fly (*k*_0_) and α = 2. Optimization is performed by taking 10,000 steps in the fitness landscape for low proliferation rates (*k*_0_ = 0.1 and 0.2) and 20,000 for the high proliferation rate (*k*_0_ = 0.5). The X axis is the number of steps taken in the fitness landscape.

**Supplemental figure 6.**
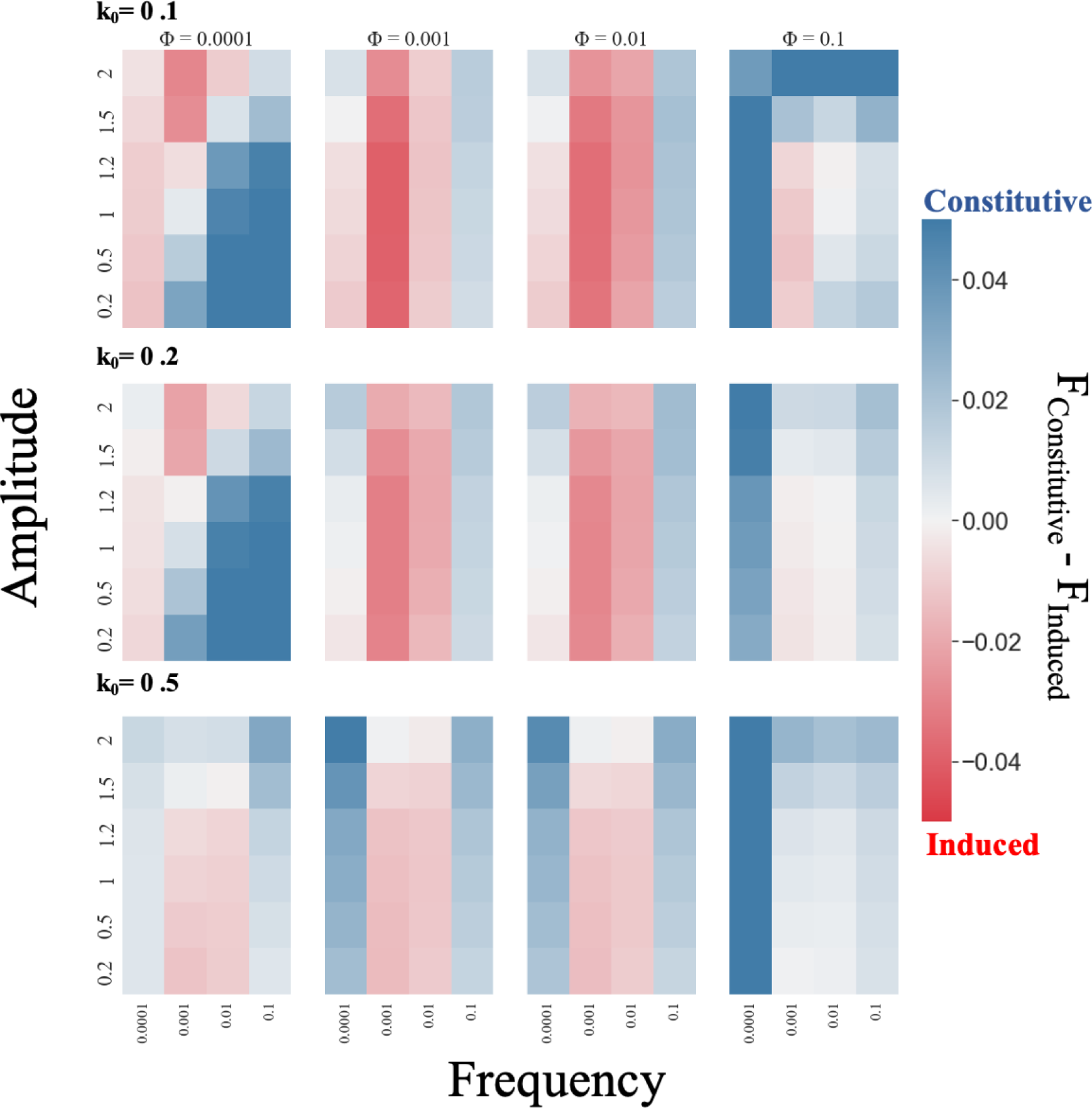
Induced responses tend to outperform constitutive defenses when flies encounter bacteria with intermediate frequencies. The relative fitness of constitutive versus induced strategies is shown as heatmaps for environments with different amplitudes and frequencies of sinusoidally oscillating encounters with bacteria. Red cells indicate that the fitness of the induced response (F_Induced_) is higher than the fitness of the constitutive response (F_Constitutive_), and blue cells indicate that F_Constitutive_ > F_Induced_. Heatmaps in the same column show induced responses that were optimized with the same frequency of the sinusoidal input of bacteria (Φ), and heatmaps in each row show the results for the same proliferation rate of bacteria (*k*_0_).

**Supplemental figure 7.**
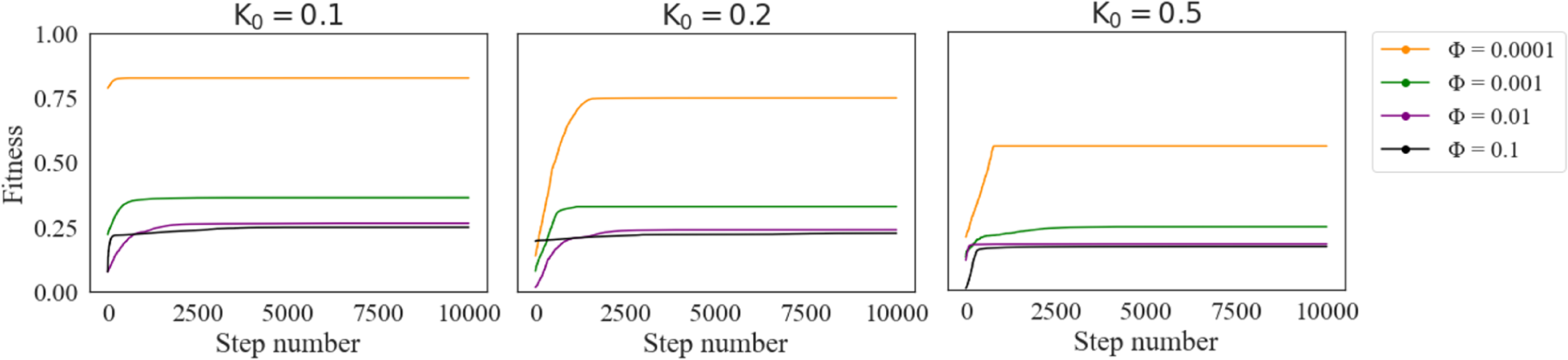
Optimization of the induced response under the assumption that production of signaling proteins does not affect the fitness. Induced defense is optimized using an oscillating deterministic input (sinusoidal) with different frequencies (Φ) for three different proliferation rates of bacteria inside the fly (*k*_0_) and α = 2. Optimization is performed by taking 10,000 steps in the fitness landscape. The X axis is the number of steps taken in the fitness landscape.

**Supplemental figure 8.**
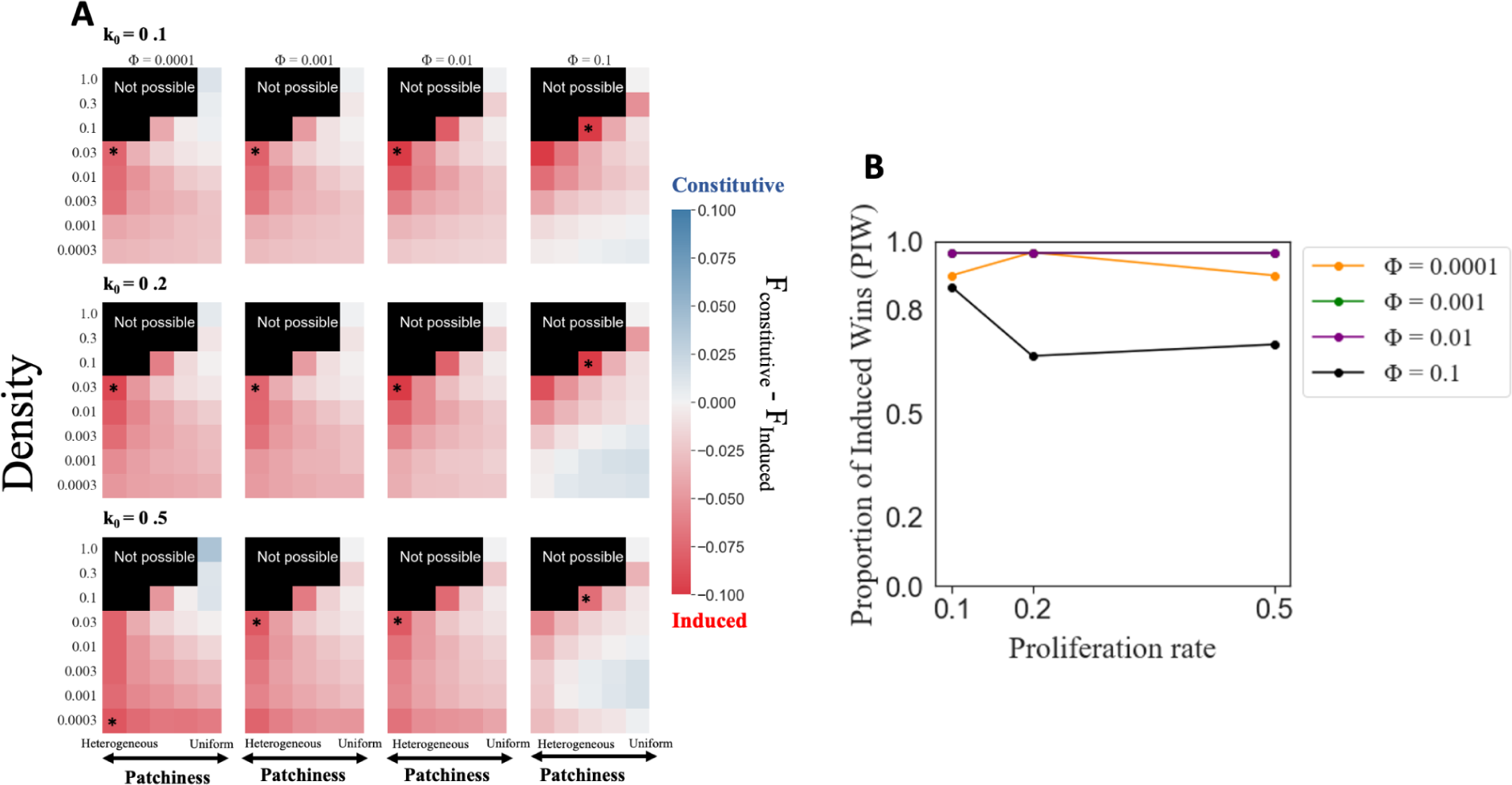
Results for comparison of constitutive and induced defenses, assuming that production of signaling proteins does not affect fitness. **A.** The relative fitness of constitutive vs induced defense (color bar) in environments with different patchiness (X-axis) and bacterial density (Y-axis). Induction has a higher fitness in red-colored cells than the best constitutive defense. Induction is preferred in heterogeneous environments with low bacterial density regardless. The highest relative fitness is shown with an asterisk. Heatmaps in the same column show induced responses that were optimized with the same frequency of the sinusoidal input of bacteria (Φ), and heatmaps in each row show the results for the same proliferation rate of bacteria (*k*_0_). **B.** The proportion of induced wins (number of red cells divided by the total number of cells) is graphed for different optimizations of the induced defense.

**Supplemental figure 9.**
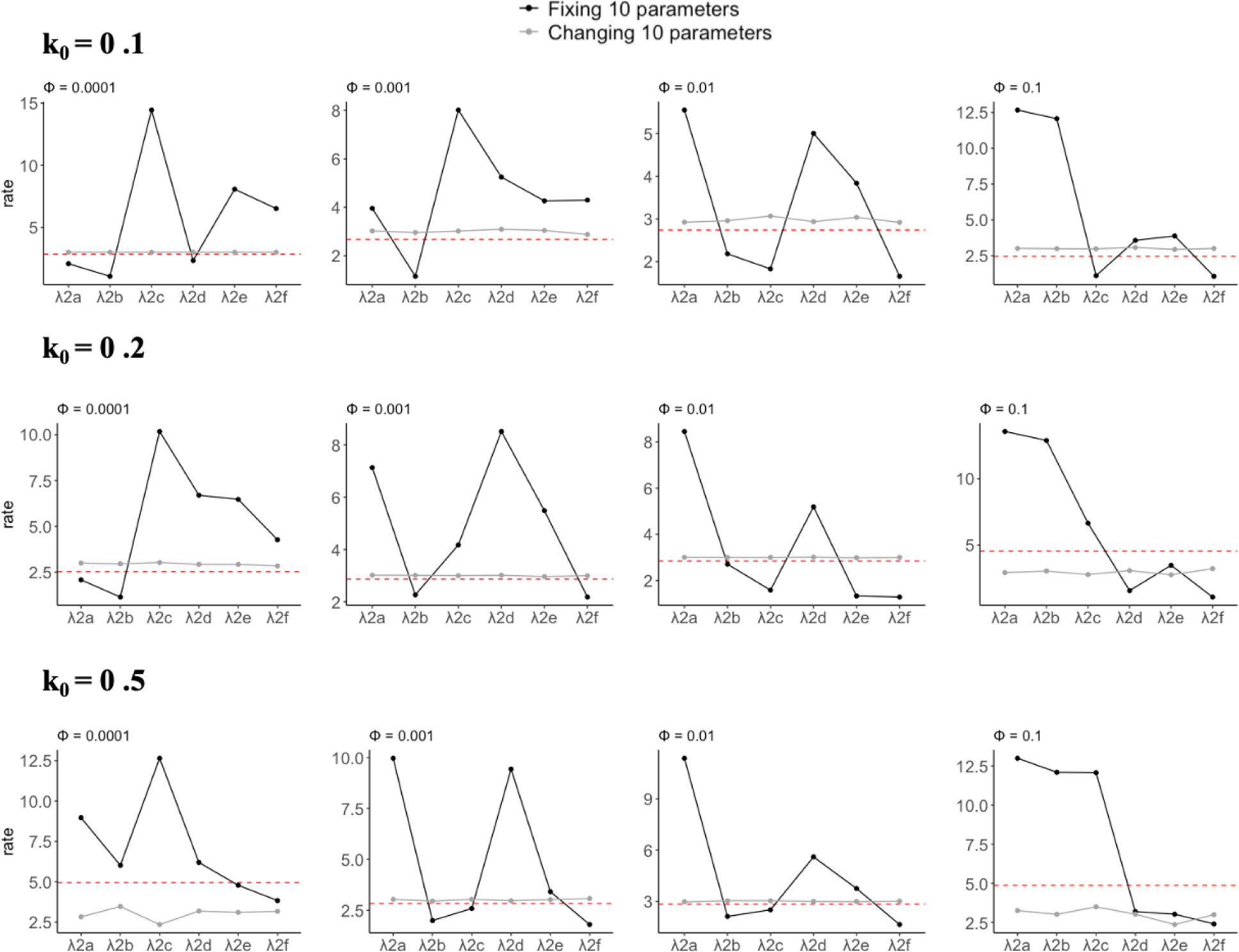
Degradation rate (Y axis) for proteins involved in Imd signaling: R (λ2_*a*_), N (λ2_*b*_), L (λ2_*c*_), P (λ2_*d*_), S (λ2_*c*_), and A (λ2_*f*_) (X-axis) are shown for two methods of optimization of the induced defense (columns) against different bacterial proliferation rates (rows). The black line shows the optimized degradation rates when the other 10 parameters are fixed during optimization. The gray line shows optimized degradation rates when all 16 parameters are allowed to fluctuate during optimization. The red dashed line is the optimized degradation rate under the assumption of identical degradation rate (λ_2_) for all proteins.

**Supplemental figure 10.**
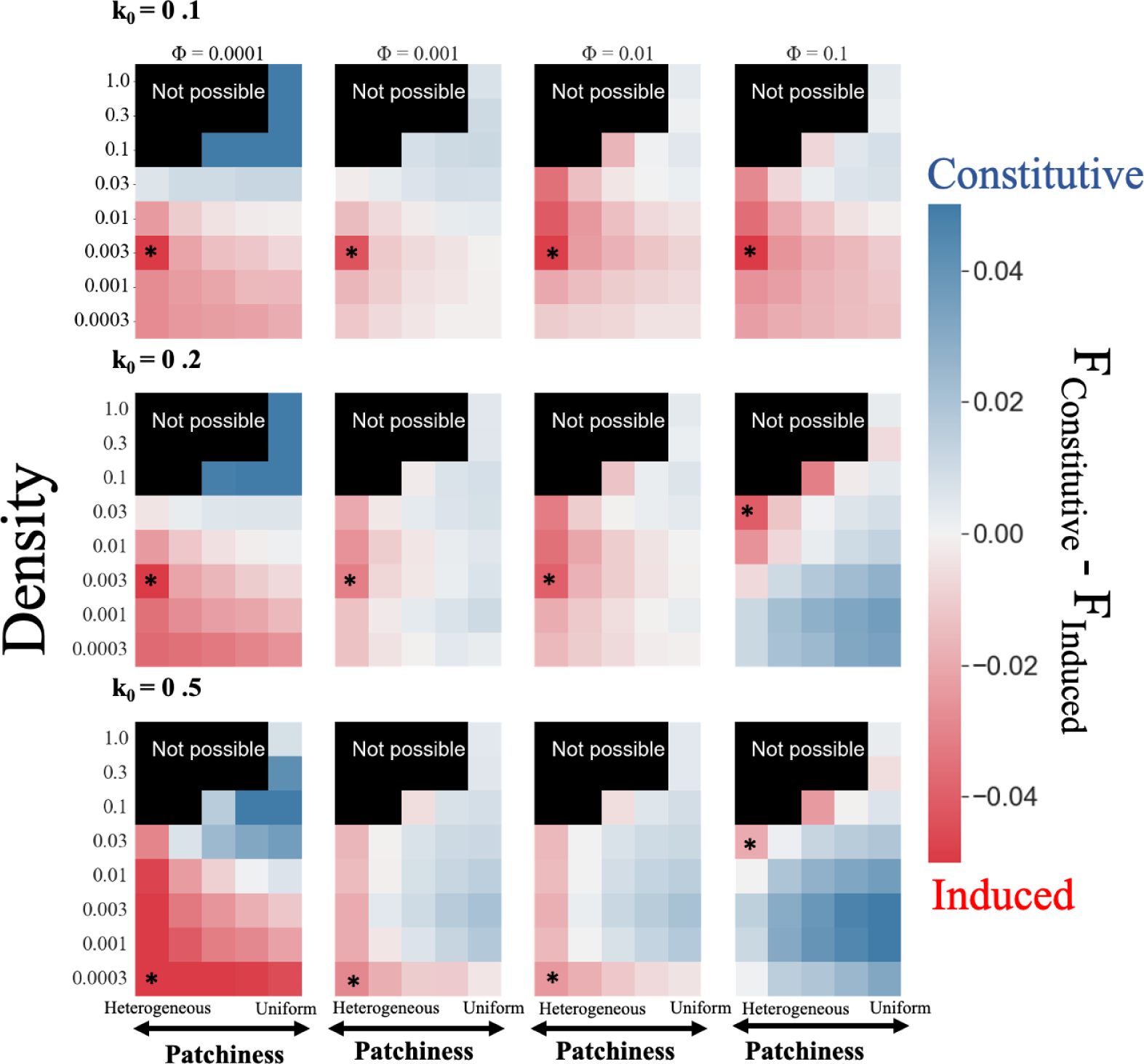
Results for comparison of constitutive and induced defenses, assuming different degradation rates for immune proteins (16-parameter model). The relative fitness of constitutive vs induced defense (color bar) in environments with different patchiness (X-axis) and bacterial density (Y-axis). Induction tends to outperform constitutive defenses when flies encounter bacteria with intermediate frequencies. Induction has a higher fitness in red-colored cells than the best constitutive defense. The highest relative fitness is shown with an asterisk. Heatmaps in the same column show induced responses that were optimized with the same frequency of the sinusoidal input of bacteria (Φ), and heatmaps in each row show the results for the same proliferation rate of bacteria (*k*_0_).

**Supplemental figure 11.**
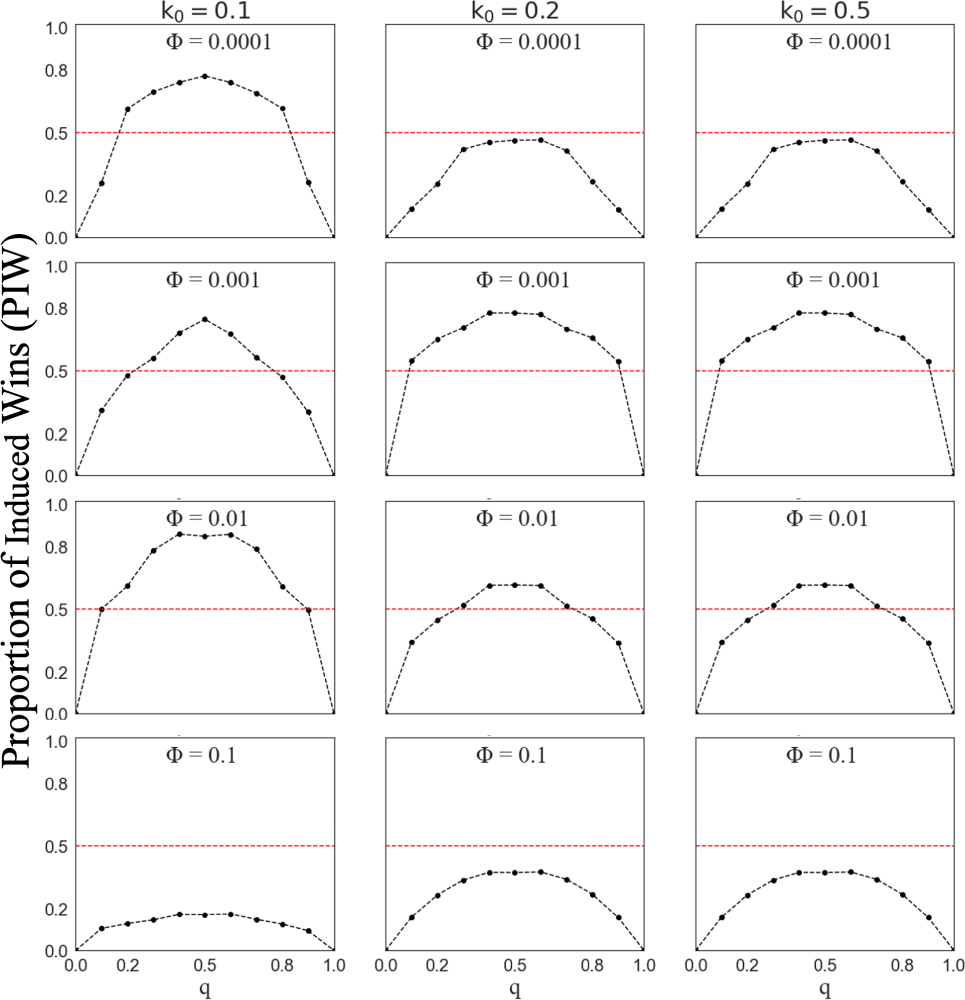
Equal frequencies in two environments favor an induced response. The proportion of times induction outperforms constitutive defense, PIW (Y-axis), is plotted against the probability of inhabiting one of the environments (X-axis). Graphs in a column have the same bacterial proliferation rate (*k*_0_), and graphs in a row have the same Φ value used to to optimize the induced response. Induction outperforms constitutive defense above the dashed line (PIW>0.5), and constitutive defense performs better below the dashed line (PIW<0.5).

**Supplemental figure 12.**
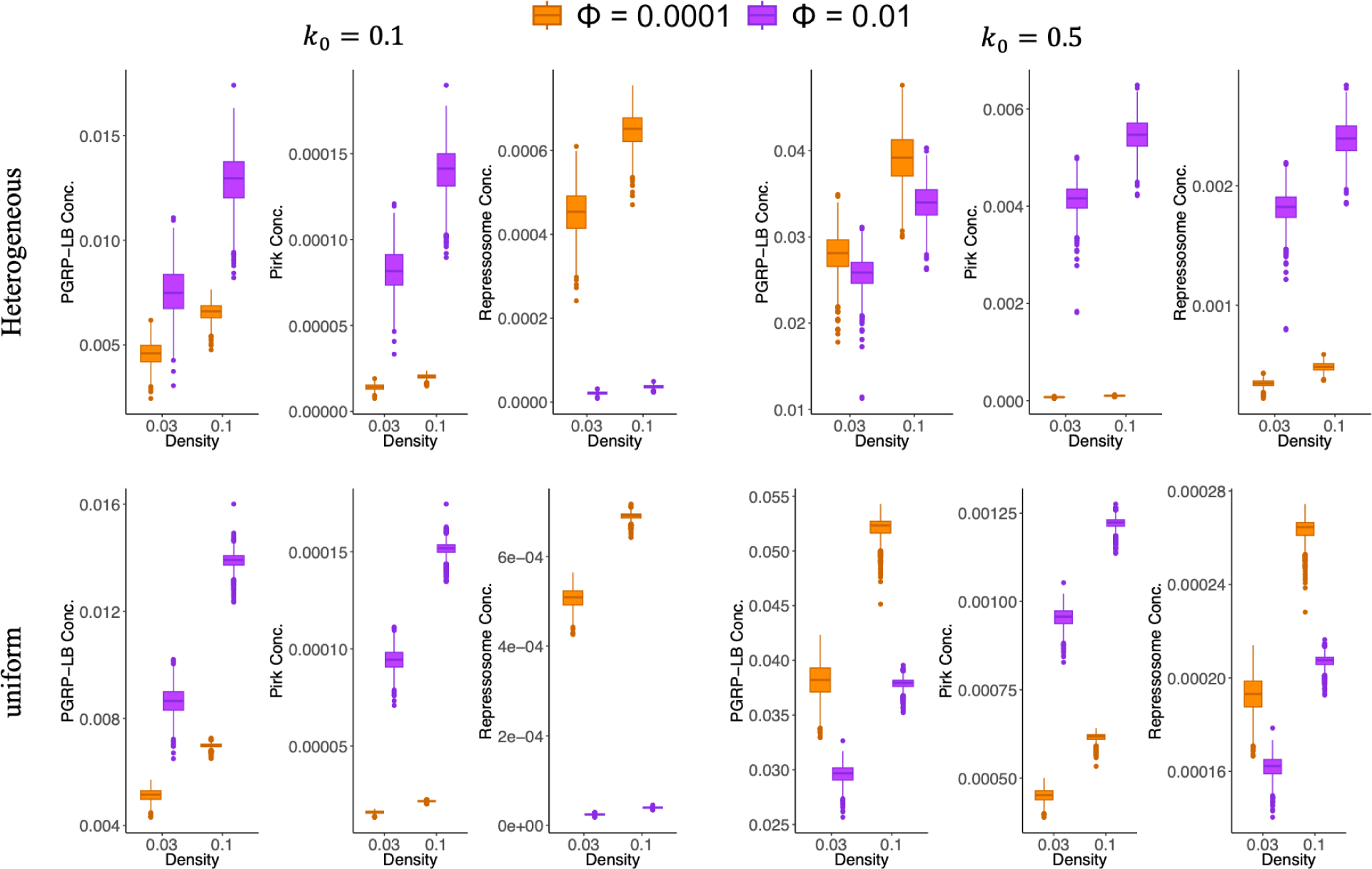
Concentration of proteins involved in negative regulation of the Imd pathway for different optimizations of the induced defense (different values of Φ) across 1,000 simulations for two bacterial densities (0.03 and 0.1). The top row shows the results for the heterogeneous (*p* = 1) distribution of bacteria and the bottom row for the uniform (*p* = 3) distribution of bacteria.

**Supplemental figure 13.**
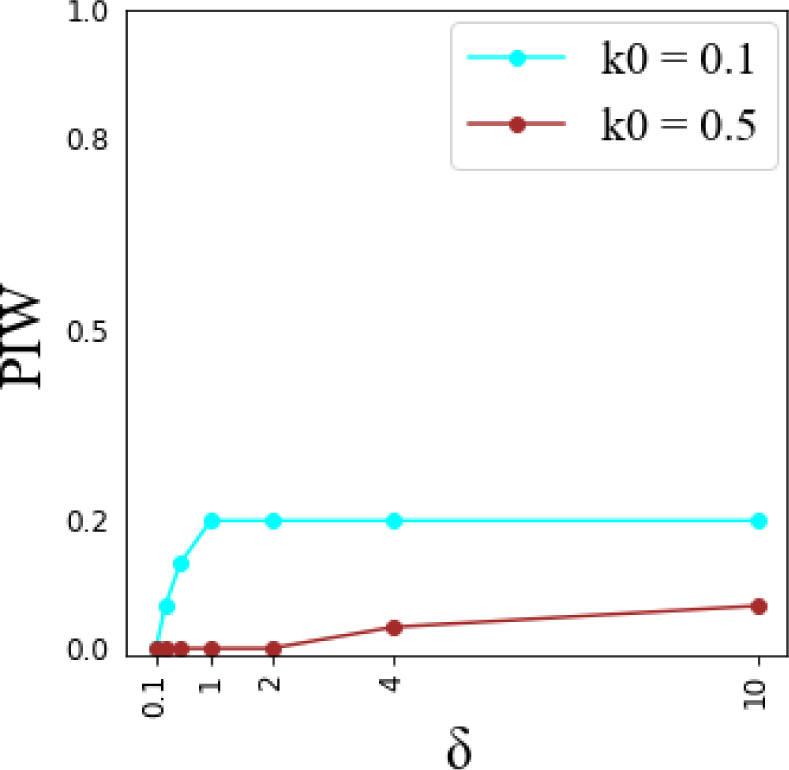
The proportion of induced wins (Y-axis) is plotted against different values of Zs (X-axis). The parameter value is changed by multiplying the optimized values for Φ = 0.1 and *k*_0_ = 0.5 by a constant (δ).

**Supplemental Table 1.**
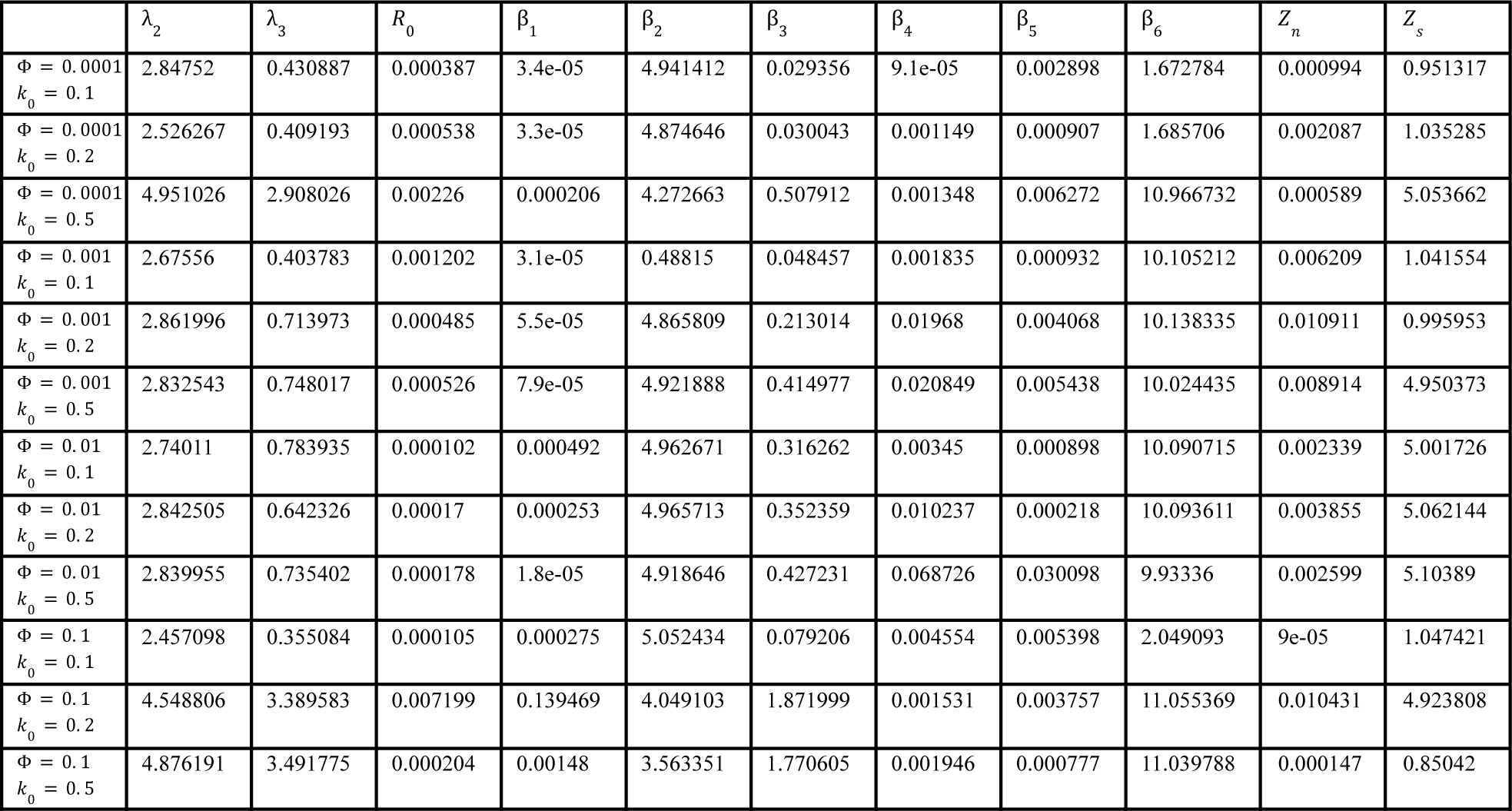
Optimized parameter values for the 11-parameter model using different frequency of exposure to bacteria (Φ) with different proliferation rates (*k*_0_).

| | $\lambda_2$ | $\lambda_3$ | $R_0$ | $\beta_1$ | $\beta_2$ | $\beta_3$ | $\beta_4$ | $\beta_5$ | $\beta_6$ | $Z_n$ | $Z_s$ |
| --- | --- | --- | --- | --- | --- | --- | --- | --- | --- | --- | --- |
| $\Phi = 0.0001$<br>$k_0 = 0.1$ | 2.84752 | 0.430887 | 0.000387 | 3.4e-05 | 4.941412 | 0.029356 | 9.1e-05 | 0.002898 | 1.672784 | 0.000994 | 0.951317 |
| $\Phi = 0.0001$<br>$k_0 = 0.2$ | 2.526267 | 0.409193 | 0.000538 | 3.3e-05 | 4.874646 | 0.030043 | 0.001149 | 0.000907 | 1.685706 | 0.002087 | 1.035285 |
| $\Phi = 0.0001$<br>$k_0 = 0.5$ | 4.951026 | 2.908026 | 0.00226 | 0.000206 | 4.272663 | 0.507912 | 0.001348 | 0.006272 | 10.966732 | 0.000589 | 5.053662 |
| $\Phi = 0.001$<br>$k_0 = 0.1$ | 2.67556 | 0.403783 | 0.001202 | 3.1e-05 | 0.48815 | 0.048457 | 0.001835 | 0.000932 | 10.105212 | 0.006209 | 1.041554 |
| $\Phi = 0.001$<br>$k_0 = 0.2$ | 2.861996 | 0.713973 | 0.000485 | 5.5e-05 | 4.865809 | 0.213014 | 0.01968 | 0.004068 | 10.138335 | 0.010911 | 0.995953 |
| $\Phi = 0.001$<br>$k_0 = 0.5$ | 2.832543 | 0.748017 | 0.000526 | 7.9e-05 | 4.921888 | 0.414977 | 0.020849 | 0.005438 | 10.024435 | 0.008914 | 4.950373 |
| $\Phi = 0.01$<br>$k_0 = 0.1$ | 2.74011 | 0.783935 | 0.000102 | 0.000492 | 4.962671 | 0.316262 | 0.00345 | 0.000898 | 10.090715 | 0.002339 | 5.001726 |
| $\Phi = 0.01$<br>$k_0 = 0.2$ | 2.842505 | 0.642326 | 0.00017 | 0.000253 | 4.965713 | 0.352359 | 0.010237 | 0.000218 | 10.093611 | 0.003855 | 5.062144 |
| $\Phi = 0.01$<br>$k_0 = 0.5$ | 2.839955 | 0.735402 | 0.000178 | 1.8e-05 | 4.918646 | 0.427231 | 0.068726 | 0.030098 | 9.93336 | 0.002599 | 5.10389 |
| $\Phi = 0.1$<br>$k_0 = 0.1$ | 2.457098 | 0.355084 | 0.000105 | 0.000275 | 5.052434 | 0.079206 | 0.004554 | 0.005398 | 2.049093 | 9e-05 | 1.047421 |
| $\Phi = 0.1$<br>$k_0 = 0.2$ | 4.548806 | 3.389583 | 0.007199 | 0.139469 | 4.049103 | 1.871999 | 0.001531 | 0.003757 | 11.055369 | 0.010431 | 4.923808 |
| $\Phi = 0.1$<br>$k_0 = 0.5$ | 4.876191 | 3.491775 | 0.000204 | 0.00148 | 3.563351 | 1.770605 | 0.001946 | 0.000777 | 11.039788 | 0.000147 | 0.85042 |

**Supplemental Table 2.** The numerical value of density (*d*) and patchiness (*p*) used to simulate bacterial population. Table B contains *d* and *p* values for cells with different cell numbers (cell#), based on table A.

**A**
|  |  |  |  |  |
| --- | --- | --- | --- | --- |
| Not possible |  |  |  | 23 |
|  |  |  |  | 24 |
|  |  | 11 | 17 | 25 |
| 1 | 6 | 12 | 18 | 26 |
| 2 | 7 | 13 | 19 | 27 |
| 3 | 8 | 14 | 20 | 28 |
| 4 | 9 | 15 | 21 | 29 |
| 5 | 10 | 16 | 22 | 30 |

**B**
| Cell # | $p$ | $d$ |
| --- | --- | --- |
| 1 | 1 | 0.03 |
| 2 | 1 | 0.01 |
| 3 | 1 | 0.003 |
| 4 | 1 | 0.001 |
| 5 | 1 | 0.0003 |
| 6 | 2 | 0.03 |
| 7 | 2 | 0.01 |
| 8 | 2 | 0.003 |
| 9 | 2 | 0.001 |
| 10 | 2 | 0.0003 |
| 11 | 1 | 0.1 |
| 12 | 3 | 0.03 |
| 13 | 3 | 0.01 |
| 14 | 3 | 0.003 |
| 15 | 3 | 0.001 |
| 16 | 3 | 0.0003 |
| 17 | 2 | 0.1 |
| 18 | 4 | 0.03 |
| 19 | 4 | 0.01 |
| 20 | 4 | 0.003 |
| 21 | 4 | 0.001 |
| 22 | 4 | 0.0003 |
| 23 | 1 | 1.0 |
| 24 | 1 | 0.3 |
| 25 | 3 | 0.1 |
| 26 | 5 | 0.03 |
| 27 | 5 | 0.01 |
| 28 | 5 | 0.003 |
| 29 | 5 | 0.001 |
| 30 | 5 | 0.0003 |

